# Dynamic Lipidome Reorganization in Response to Heat Shock Stress

**DOI:** 10.1101/2025.02.18.638884

**Authors:** Luis Solano, Uri Keshet, Andrew Reinschmidt, Yonny Chavez, William Drew Hulsy, Oliver Fiehn, Nikolas Nikolaidis

## Abstract

The heat shock response (HSR) is a conserved cellular mechanism critical for adaptation to environmental and physiological stressors, with broad implications for cell survival, immune responses, and cancer biology. While the HSR has been extensively studied at the proteomic and transcriptomic levels, the role of lipid metabolism and membrane reorganization remains underexplored. Here, we integrate mass spectrometry-based lipidomics with RNA sequencing to characterize global lipidomic and transcriptomic changes in HeLa cells exposed to three conditions: control, heat shock (HS), and HS with eight hours of recovery. Heat shock-induced extensive lipid remodeling, including significant increases in fatty acids, glycerophospholipids, and sphingolipids, with partial normalization during recovery. Transcriptomic analysis identified over 2,700 upregulated and 2,300 downregulated genes under heat shock, with GO enrichment suggesting potential transcriptional contributions to lipid metabolism. However, transcriptional changes alone did not fully explain the observed lipidomic shifts, suggesting additional layers of regulation. Joint pathway analysis revealed enrichment in glycerophospholipid and sphingolipid metabolism, while network analysis identified lipid transport regulators (STAB2, APOB), stress-linked metabolic nodes (KNG1), and persistent sphingolipid enrichment during recovery. These findings provide a comprehensive framework for understanding lipid-mediated mechanisms of the HSR and highlight the importance of multi-omics integration in stress adaptation and disease biology.

## 1. Introduction

The heat shock response (HSR) is a conserved cellular mechanism enabling cells to adapt to environmental and physiological stressors, such as elevated temperatures, oxidative stress, and metabolic challenges [1,2]. This response is classically characterized by the induction of heat shock proteins (HSPs), which act as molecular chaperones to maintain protein homeostasis [3,4]. However, beyond protein regulation, the HSR encompasses broader cellular reprogramming, including significant changes in lipid metabolism and membrane composition [5-7]. These lipidomic changes are critical for maintaining membrane integrity, cellular signaling, and stress adaptation [8-10].

Lipids play a central role in cellular stress responses by contributing to membrane remodeling, intracellular signaling, and energy storage [11-13]. Heat stress induces profound lipidomic changes, including increased levels of fatty acids, sphingolipids, and phosphatidylserine (PS) [14,15]. These changes alter the biophysical properties of the plasma membrane (PM), such as fluidity and rigidity, which are essential for stress sensing and cellular adaptation [8-10]. For example, elevated sphingolipids and ceramides have been implicated in membrane stabilization during stress, while increases in PS facilitate signaling pathways critical for the stress response [12]. Additionally, changes in cholesterol levels and phosphatidylethanolamine (PE) influence the organization of lipid rafts, which serve as hubs for signal transduction [16].

Despite the well-characterized role of transcriptional regulation in the heat shock response, how lipid remodeling is regulated at the molecular level remains unclear [17-20]. Prior studies suggest that transcriptional changes alone may not fully account for lipidomic shifts during heat stress, indicating that alternative regulatory mechanisms, such as enzyme activity and lipid trafficking, may be at play [18,21]. Emerging evidence suggests that these lipidomic shifts may not be primarily driven by transcriptional regulation but by enzymatic activity, metabolic flux adjustments, and post-transcriptional modifications [22-25]. This evidence underscores the need to investigate non-genomic mechanisms governing lipid metabolism in stress adaptation [19,26].

Across biological systems, lipid remodeling is a conserved adaptive strategy for coping with thermal and metabolic stress [18,21-25]. In plants, bacteria, and mammalian cells, membrane composition, and lipid biosynthesis shifts support stress tolerance by modulating membrane fluidity, vesicle trafficking, and energy homeostasis [21,27-30]. Similar lipidomic shifts are observed in cancer cells, where altered lipid metabolism is a hallmark of malignancy [31-33]. Cancer cells reprogram lipid biosynthesis to sustain rapid proliferation and survive in hostile microenvironments, such as hypoxia and oxidative stress [31,34,35]. For instance, increased sphingolipid metabolism and fatty acid synthesis are associated with enhanced cell survival and metastasis, while altered phosphatidylcholine (PC) to PE ratios disrupt membrane homeostasis and signaling [33,36]. The parallels between lipid remodeling during the HSR and in cancer suggest that these processes share overlapping mechanisms, highlighting the importance of lipids in both physiological and pathological stress responses [37,38].

In this study, we integrate mass spectrometry-based lipidomics with transcriptomic analyses to dissect the regulation of lipid remodeling in the heat shock response. Because lipid metabolism may be primarily regulated at the enzymatic and post-transcriptional level [39], this multi-omics approach provides a comprehensive framework for understanding the metabolic adjustments that enable cells to withstand thermal stress [40,41]. Our study focuses on identifying key pathways and molecular processes driving lipidome reorganization. By addressing the critical gap in understanding how lipid metabolism contributes to the HSR, we aim to elucidate mechanisms that could inform therapeutic strategies for diseases linked to stress adaptation, including cancer.

## 2. Results

### 2.1. Global Lipidomic Changes in Response to Heat Shock

#### Principal Component Analysis (PCA) Reveals Distinct Lipidomic Profiles

Principal component analysis (PCA) was performed to assess the global variance in lipid composition across control, heat shock (R0), and recovery (R8) conditions. The scree plot (Figure 1A) demonstrates that the first three principal components explain more than 95% of the total variance, with PC1 capturing the majority of separation between experimental conditions. The PCA score plot (Figure 1B) shows an apparent clustering of lipid profiles based on treatment, with R0 samples diverging significantly from control along PC1. Interestingly, R8 samples shift away from R0 and toward the control cluster, indicating a partial lipidome recovery after 8 hours. These results confirm that heat shock induces a widespread reorganization of the lipidome, which begins to normalize during recovery.

**Figure 1.**
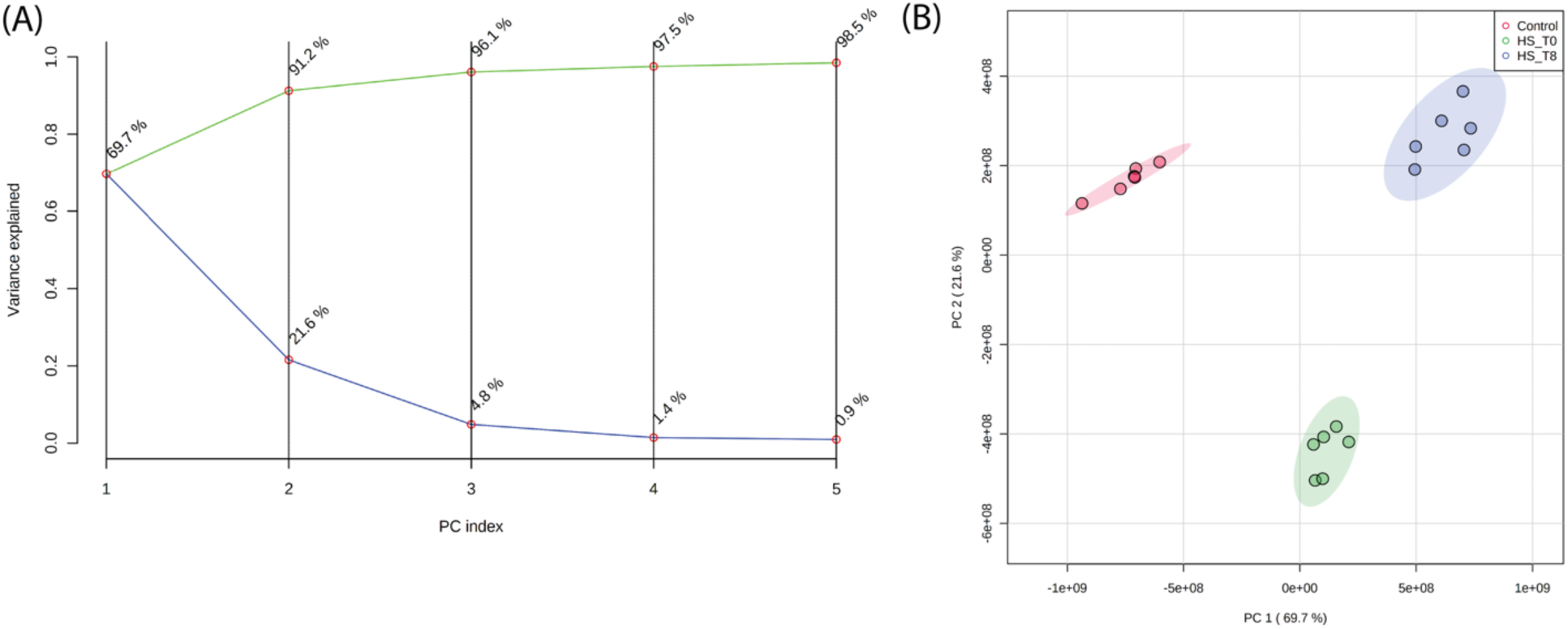
Principal Component Analysis (PCA) of Lipidomic Profiles reveals distinct clustering of lipidomic profiles across experimental conditions. (A) A scree plot shows that the first three principal components explain >95% of the total variance. (B) PCA score plot demonstrating distinct clustering of lipidomic profiles based on experimental conditions. PC1 primarily separates heat shock (R0) from control and recovery (R8), with R8 samples shifting closer to control, indicating partial lipidome normalization.

#### ANOVA Identifies Lipids Significantly Altered by Heat Shock

A one-way ANOVA followed by Fisher’s least significant difference post hoc test identified 771 lipids with significant abundance changes (p < 0.05) across experimental conditions (Figure 2). Most significantly altered lipids were observed in R0 vs. control, suggesting that the immediate heat shock response involves substantial lipid remodeling. The number of differentially abundant lipids remains high in R8 vs. R0 but with a distinct subset exhibiting sustained elevation or return to baseline levels, suggesting dynamic regulation of lipid metabolism throughout the stress-recovery process.

**Figure 2.**
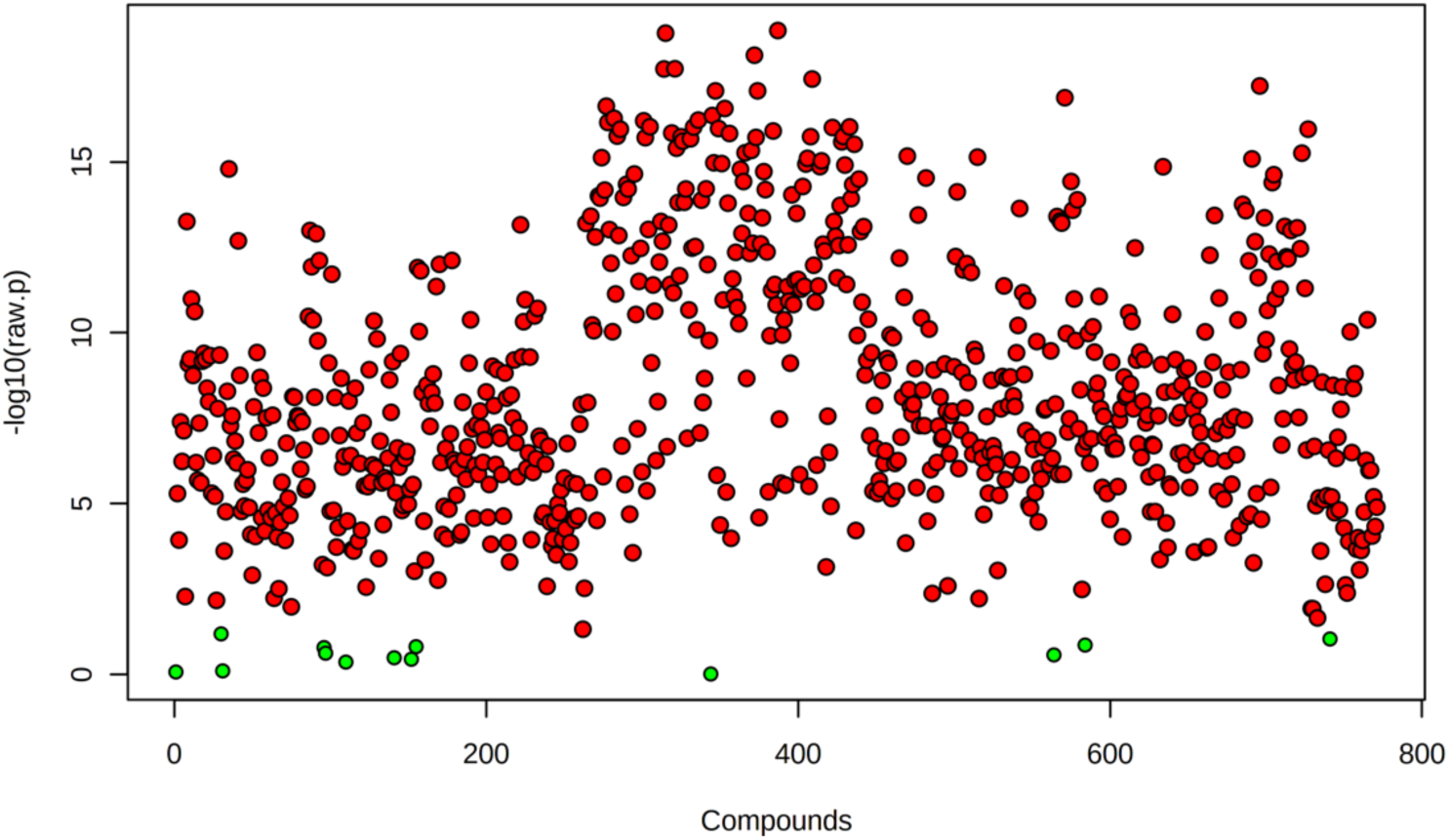
Heat Shock significantly alters the lipidome. Volcano plot displaying 771 lipids with significant abundance changes (One-way ANOVA; *P* < 0.05) across experimental conditions. Each dot represents an individual lipid, with red indicating significantly altered lipids identified through Fisher’s Least Significant Difference (LSD) post hoc test, while green dots represent non-significant changes. The y-axis (-log10(*P-*value)) indicates the statistical significance of lipid changes, with higher values reflecting more substantial evidence for differential abundance. The x-axis represents different lipid species arranged arbitrarily.

#### General Shifts in Major Lipid Classes

Total lipid abundance was quantified across conditions (Figure 3A, B) to evaluate broad lipidomic changes. Heat shock resulted in a 35% increase in total lipid content compared to control, primarily driven by changes in phospholipids, sphingolipids, and free fatty acids. By R8, total lipid abundance showed a partial reduction, suggesting activation of lipid turnover mechanisms during recovery.

**Figure 3.**
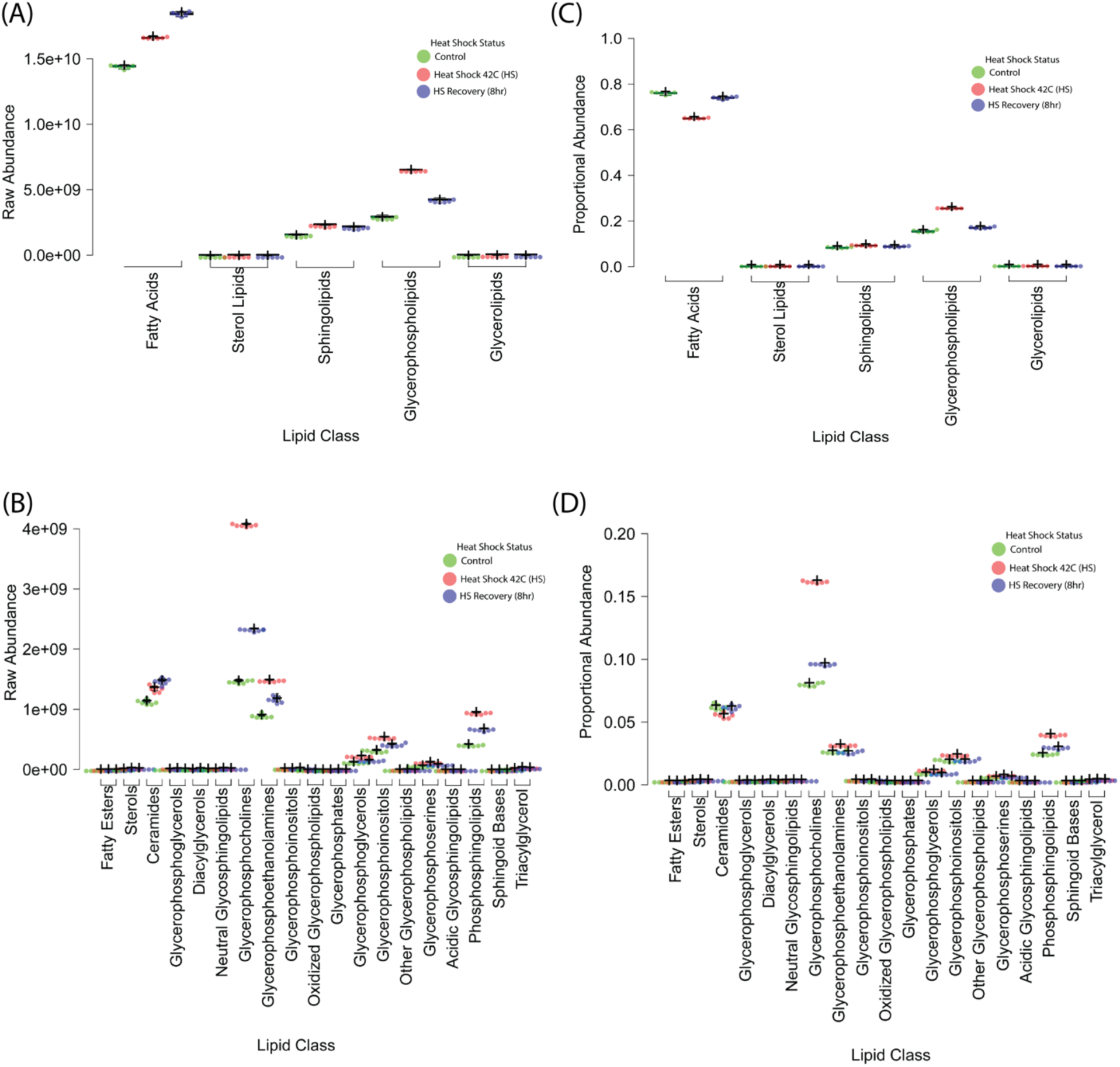
Global Lipidomic Changes in Response to Heat Shock. (A, B) Total lipid abundance increased by 35% upon heat shock (R0), with a partial reduction at R8, indicating lipid turnover during recovery. Phospholipids, sphingolipids, and free fatty acids showed the most pronounced changes, while sterol lipids remained relatively stable. (C, D) Lipid class composition shifts in response to heat shock and recovery. Phospholipid subclasses (PS, PE, PG, PI, PC), sphingolipids, and ceramides significantly increased during heat shock, supporting membrane remodeling and stress adaptation. Free fatty acids (FA) increased sharply at R0 and continued rising in recovery (R8), suggesting sustained metabolic activity. Sterol lipids remained essentially unchanged. Boxplots display median (center lines), 25th and 75th percentiles (box limits), and whiskers extending 1.5 times the interquartile range. Dots represent outliers, crosses indicate sample means, and bars represent 95% confidence intervals (n = 6 biological replicates).

Proportional composition analysis (Figure 3C, D) revealed that membrane-associated lipid classes exhibited the highest shifts, and phospholipid subclasses and sphingolipids increased significantly in response to heat shock. These shifts suggest that membrane remodeling is a key feature of the heat shock response, potentially altering biophysical properties such as fluidity and curvature.

#### Heatmaps Reveal Distinct Lipidomic Clusters Across Conditions

Hierarchical clustering heatmaps were generated to visualize further the lipidomic shifts across conditions (Figure 4 and Supplementary Figure S1). These heatmaps demonstrate the apparent clustering of samples based on lipid composition, aligning with PCA results (Figure 1B). The R0 condition exhibits a striking increase in specific lipid subclasses, while R8 samples display a partial return toward the control profile. Notably, clusters of phospholipids and sphingolipids showed persistent elevation in R8, suggesting a prolonged role in membrane remodeling and stress recovery. These heatmap findings support the PCA and ANOVA results, confirming that lipidomic remodeling during heat shock follows a dynamic, condition-dependent pattern

**Figure 4.**
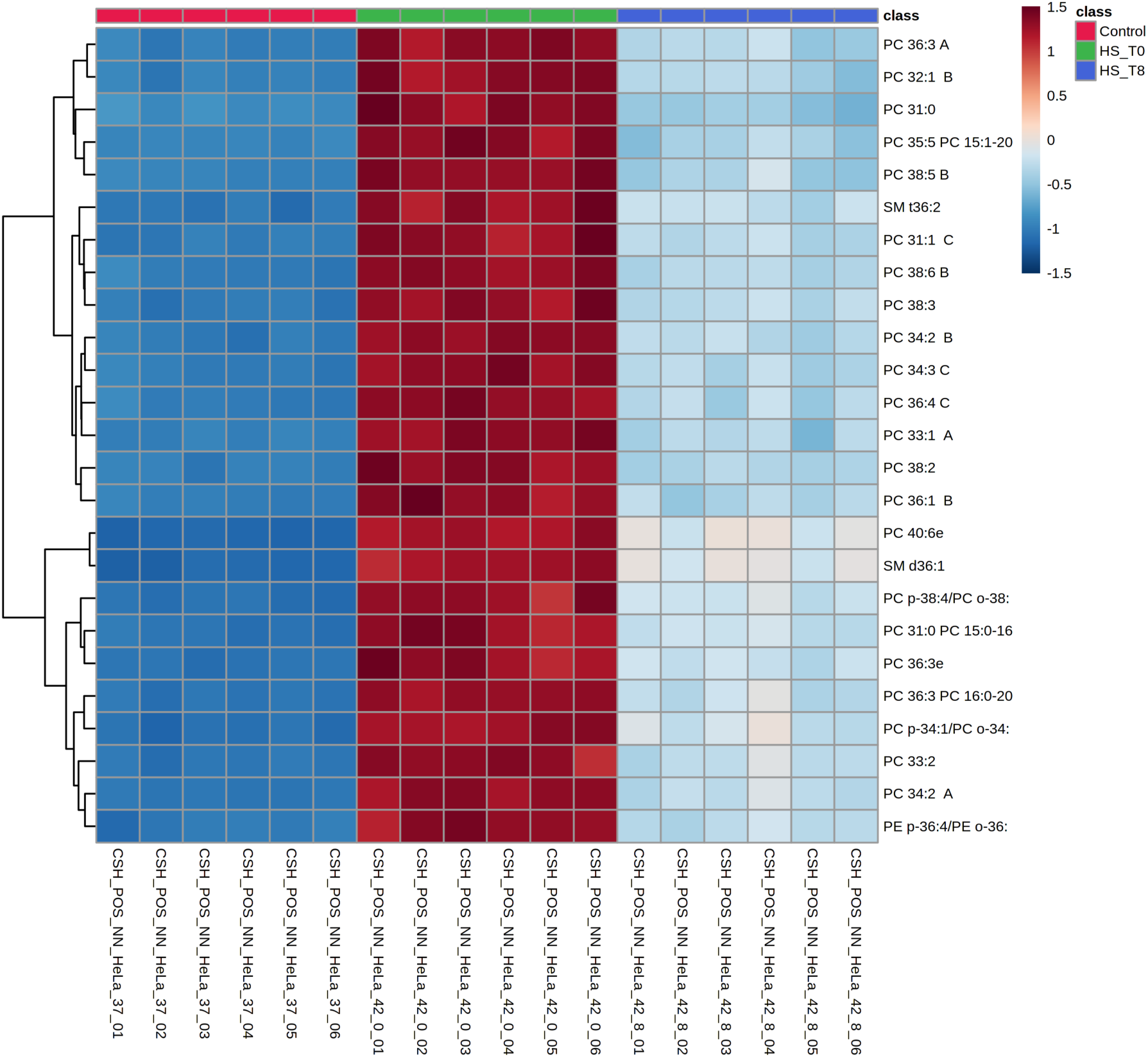
Hierarchical Clustering of Lipidomic Profiles demonstrates that PC species dominate the top 25 differentially abundant lipids, implying dynamic lipidome remodeling in response to heat shock. Heatmap depicting hierarchical clustering of the top 25 differentially abundant lipids (One-way ANOVA; *P* < 0.05) across experimental conditions. Lipid intensities are represented as Z-scores, with blue indicating decreased abundance and red indicating increased abundance relative to the mean. Experimental groups are color-coded at the top: control (green), HS_T0 (red; heat shock), and HS_T8 (blue; recovery at 8 hours post-heat shock). Lipid subclasses such as phosphatidylcholines (PCs), sphingomyelins (SMs), and phosphatidylethanolamines (PEs) exhibit significant shifts upon heat shock. Some lipids remain partially elevated at R8, suggesting prolonged lipid remodeling and delayed recovery dynamics. The clustering pattern highlights lipidomic reorganization as a critical component of the heat shock response.

#### Paired Univariate Statistical Analysis Highlights Key Lipid Changes

Volcano plots from paired univariate statistical analysis (Figure 5 and Supplementary Figure S2) revealed specific lipid species with significant fold changes across conditions. Paired univariate statistical analyses demonstrated that over 70% of the lipidome exhibited significant abundance shifts under heat shock (R0 vs. control), with a subset maintaining altered levels during recovery (R8 vs. R0).

**Figure 5.**
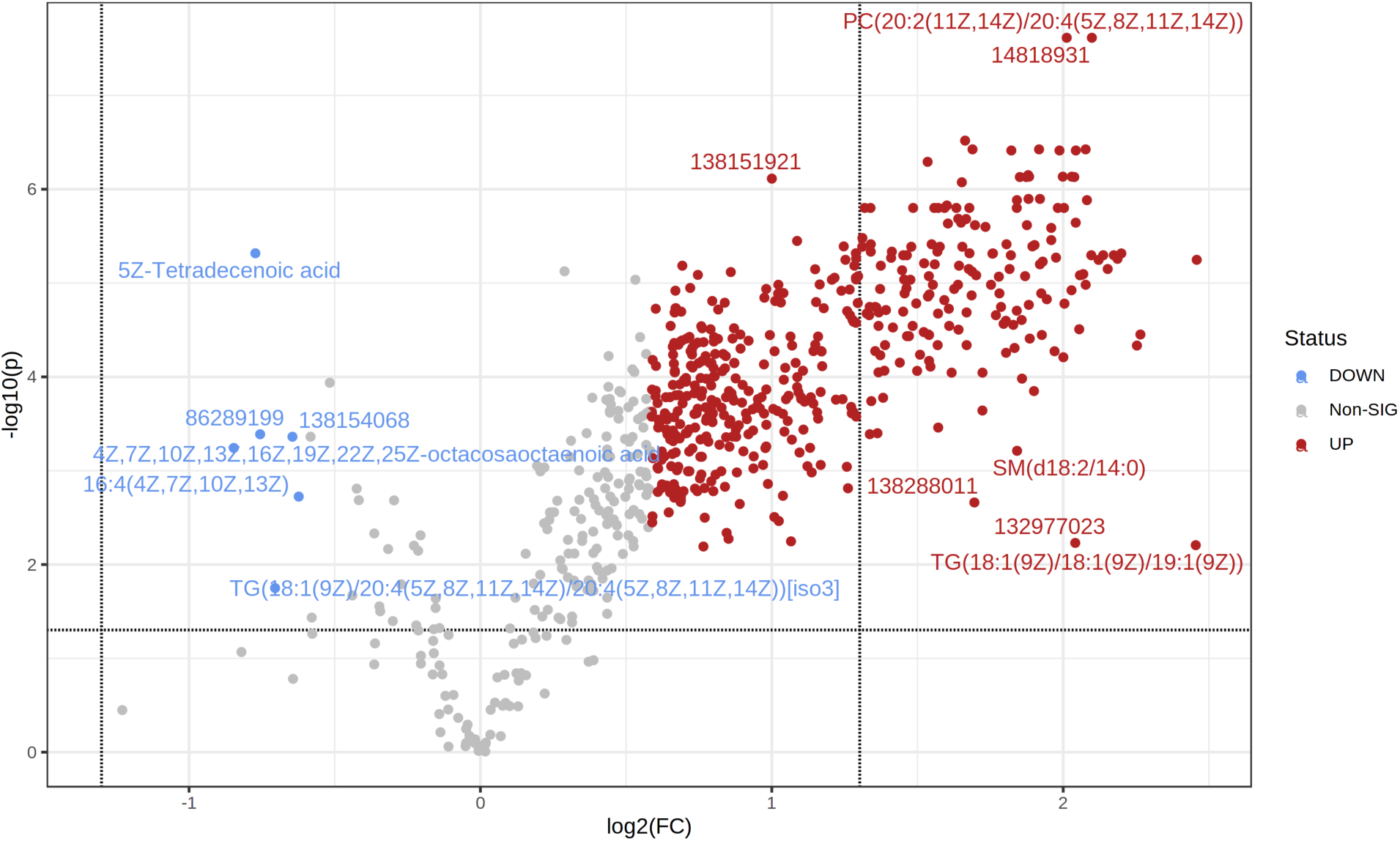
Differential Lipid Abundance in Heat Shock and Recovery highlights major lipidomic shifts post-stress. Volcano plot displaying the distribution of differentially abundant lipids in the R0 vs. control comparison. More than 70% of the lipidome exhibits significant changes upon heat shock. Lipids with increased abundance following heat shock are shown in red, whereas lipids with decreased abundance are in blue. Lipids in grey either fail to meet the fold-change cutoff (|FC| > 1.5) or do not reach statistical significance (*P*. adj < 0.05). Selected lipids of interest are labeled with their PubChem Compound Identifiers (CID), highlighting major lipid species undergoing rapid reorganization in response to heat stress.

Specifically, In the R0 vs. control comparison, 494 lipids were significantly upregulated, while 213 lipids were downregulated. For the R8 vs. control comparison, 268 lipids showed significant upregulation, reflecting sustained metabolic adaptation even during recovery. The R8 vs. R0 comparison identified 40 upregulated and 213 downregulated lipids, demonstrating a transition from stress to recovery states. These results confirm that lipid regulation during heat shock and recovery is highly dynamic, with specific lipids playing key roles in each phase.

### 2.2. Specific Lipid Alterations and Pathway Enrichment

#### Characterization of Lipid Subclasses and Their Heat Shock Response

Individual lipid subclasses were analyzed for specific trends in abundance changes to further resolve the nature of lipidomic alterations under heat shock. A breakdown of lipid categories (Figure 3 and Supplementary Figure S1) revealed significant shifts across phospholipids, sphingolipids, and free fatty acids, highlighting their potential roles in stress adaptation.

#### Fatty Acid and Phospholipid Dynamics

Heat shock resulted in a substantial increase in free fatty acids (Figure 3), with the most pronounced changes observed in unsaturated species, suggesting their involvement in membrane fluidity modulation and stress signaling. Unsaturated fatty acids are known to increase membrane flexibility, which could facilitate stress-induced remodeling of the plasma membrane [8-10].

Among phospholipids, phosphatidylcholine (PC), phosphatidylserine (PS), and phosphatidylethanolamine (PE) were the most dynamically altered lipids, exhibiting significant enrichment during heat stress, followed by partial normalization during recovery (Figures 3, 4 and Supplementary Figure S1). The increase in PS and PE is particularly relevant, as these lipids are known regulators of membrane curvature and vesicular trafficking, which may aid in the relocalization of stress-associated proteins during heat shock [8-10]. The sustained elevation of these phospholipids in R8 suggests lipid-driven membrane remodeling may persist beyond the immediate stress phase, contributing to cellular adaptation.

#### Sphingolipid Remodeling Under Heat Stress

Sphingolipids, particularly ceramides and sphingomyelins, exhibited marked increases following heat shock (Figures 3 and 5). Sphingomyelin levels peaked during heat stress but remained elevated during recovery, suggesting a sustained role in membrane stability and stress signaling. Increased ceramide abundance is particularly noteworthy, as ceramides are key regulators of apoptotic pathways and cellular stress responses. Their accumulation suggests that, under extreme stress conditions, sphingolipid metabolism may contribute to both protective adaptation and programmed cell death mechanisms.

These lipidomic alterations reveal that heat-induced stress triggers a coordinated remodeling of membrane lipids involving structural (phospholipid) and signaling (sphingolipid) changes. The observed shifts in lipid composition are likely to affect membrane biophysical properties, such as fluidity, curvature, and protein-lipid interactions, facilitating the activation of downstream stress-response pathways [42].

#### Pathway Enrichment Analysis

Pathway enrichment analysis (Figure 6, Supplementary Figure S3, and Supplementary Table S1) provided further insight into the biological processes driving lipidomic alterations. Over-representation analysis using KEGG and SMPDB databases identified key metabolic pathways impacted by heat stress. However, after correction for multiple comparisons, most pathways did not reach statistical significance. Though not statistically significant, the most enriched pathways included glycerophospholipid metabolism, sphingolipid metabolism, and biosynthesis of unsaturated fatty acids. Glycerophospholipid metabolism was enriched due to the widespread remodeling of membrane lipids, consistent with the observed increases in PS and PE, which play a role in stress adaptation and vesicle trafficking. Sphingolipid metabolism was also among the most affected pathways, with increased ceramide and sphingomyelin levels suggesting a role in stress signaling and membrane stability. Additionally, biosynthesis of unsaturated fatty acids appeared enriched, aligning with the elevated levels of free fatty acids following heat shock.

**Figure 6.**
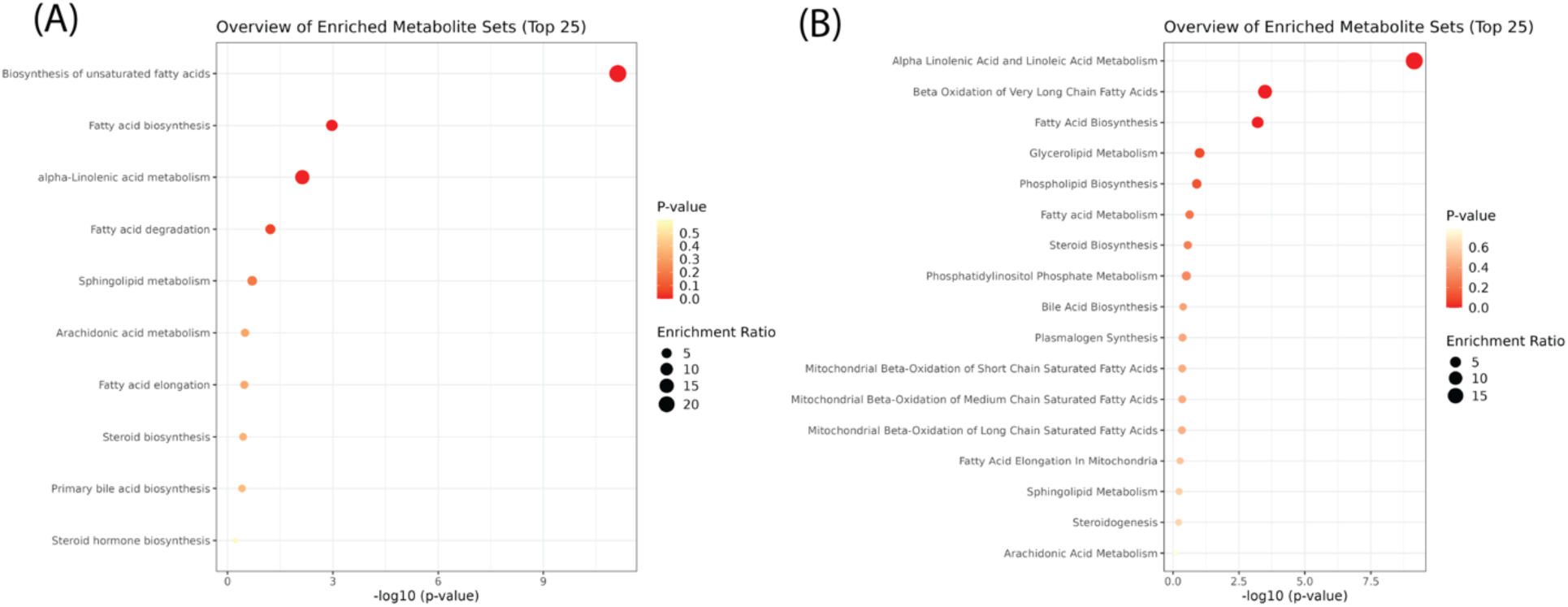
Pathway Enrichment Analysis of Lipids after Heat Shock. Over-representation analysis using (A) KEGG and (B) SMPDB databases revealed no statistically significant metabolic pathway enrichment after post hoc corrections. The x-axis shows -log10(*P*-value), where higher values indicate greater statistical significance. The y-axis lists the enriched pathways, with a focus on lipid metabolism. Each dot represents a metabolite set, colored by *P*-value (red = more significant, yellow = less significant). Dot size represents the enrichment ratio, with larger dots indicating greater pathway involvement.

These findings establish a clear metabolic trajectory wherein lipid remodeling during heat stress is driven by distinct but dispersed molecular changes rather than a single dominant pathway. The following section will explore transcriptomic responses to heat shock to elucidate further the regulatory networks orchestrating these lipidomic changes.

### 2.3. Transcriptomic Responses to Heat Shock

#### Principal Component Analysis (PCA) of Transcriptomic Changes

Principal component analysis (PCA) was performed to assess global variance in gene expression across control, heat shock (R0), and recovery (R8) conditions. The scree plot (Figure 7A) indicates that the first five principal components explain the most variance, with PC1 (43.8%) capturing the most considerable differences between conditions. The PCA score plot (Figure 7B) shows an apparent clustering of experimental groups, with R0 samples diverging significantly from the control along PC1, while R8 samples shift closer to the control, suggesting transcriptional recovery.

**Figure 7.**
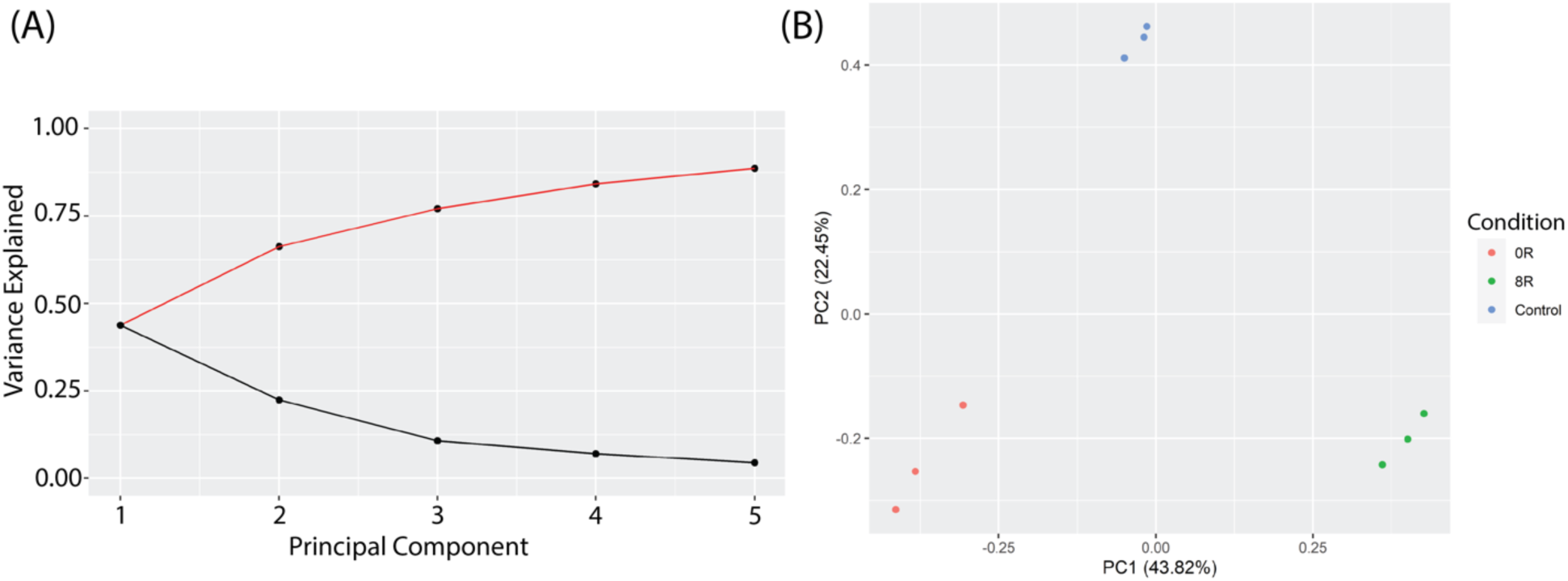
Principal Component Analysis (PCA) of Transcriptomic Profiles in Response to Heat Shock. (A) A scree plot displays the proportion of variance explained by the first five principal components (PCs). The first two PCs capture the majority of transcriptomic variance, with PC1 contributing the most. (B) PCA score plot showing distinct clustering of experimental groups. R0 (red) samples diverge significantly from control (blue) along PC1 (43.32% variance explained), indicating a strong immediate transcriptional response to heat shock. R8 (green) samples shift back toward the control group along PC1 and PC2 (22.45% variance explained), suggesting a partial recovery of transcriptomic profiles after 8 hours.

#### Differential Gene Expression Analysis

Using DESeq2, we identified differentially expressed genes (DEGs) across experimental comparisons. In the R0 vs. control condition, 2,729 genes were upregulated, while 2,377 genes were downregulated (adjusted *P-*value < 0.05). Volcano plot analysis further highlighted 1,371 significantly upregulated genes and 434 downregulated considerably genes (|log₂FC| > 1.5, adjusted *P-*value < 0.05) (Figure 8). In the R8 vs. control condition, 2,183 genes were upregulated, while 2,264 genes were downregulated, with a subset of 543 upregulated and 403 downregulated genes meeting stringent significance thresholds (Supplementary Figure S4A). The R8 vs. R0 comparison exhibited the most pronounced transcriptomic shifts, with 4,194 genes upregulated and 4,154 genes downregulated, including 569 genes with significant upregulation and 1,420 genes with significant downregulation (Supplementary Figure S4B). These results indicate a large-scale transcriptional reorganization in response to heat shock, followed by a partial recovery phase in R8.

**Figure 8.**
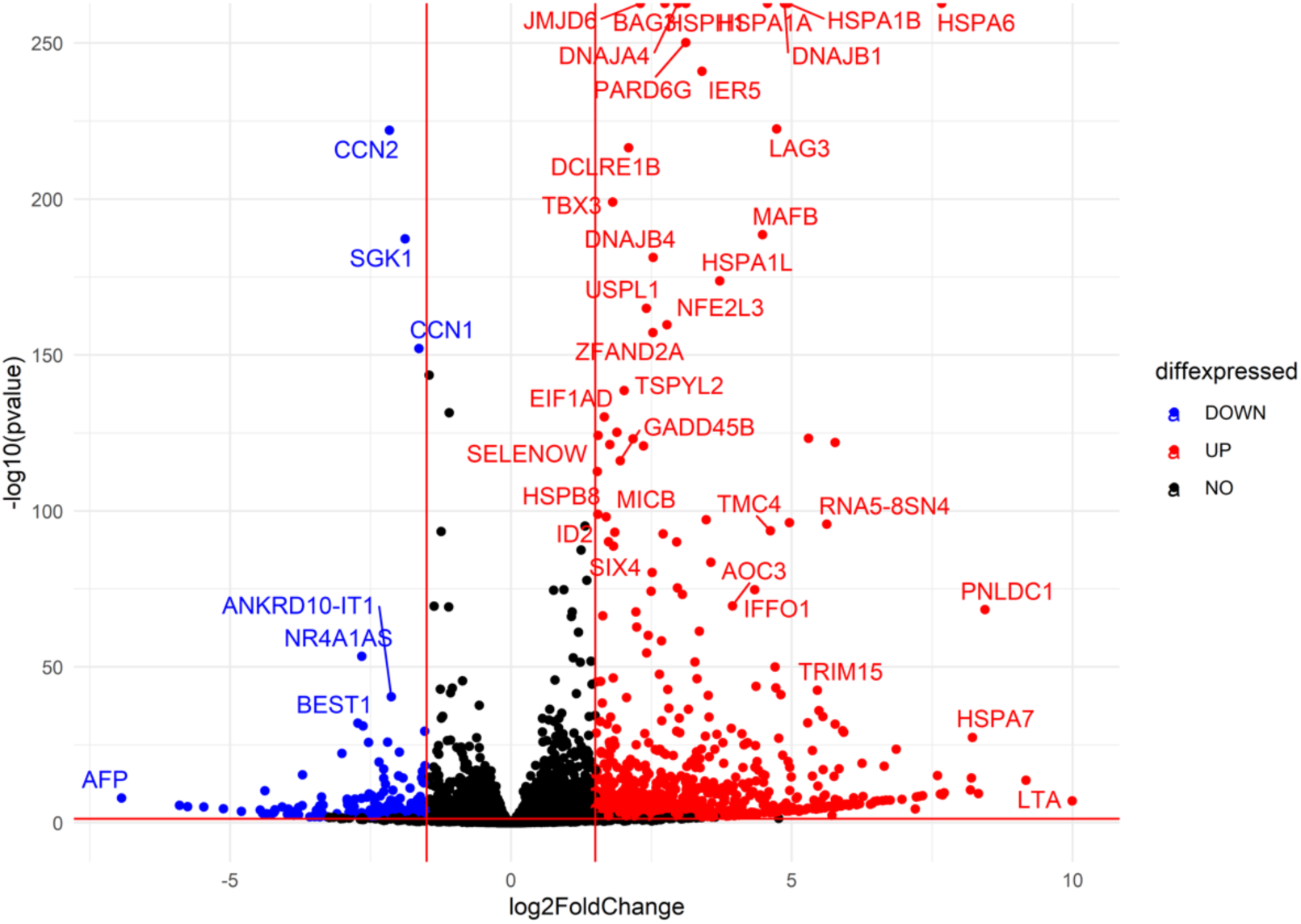
Differential Gene Expression Analysis identifies key upregulated heat shock genes and stress-responsive factors. Volcano plot depicting differentially expressed genes in R0 vs. control. Each point represents a single gene, plotted by log2(Fold Change) on the x-axis and -log10(*P*-value) on the y-axis. Upregulated genes (red): Genes significantly increased in expression upon heat shock (log2FC > 1.5, adjusted *P*-value < 0.05). Heat shock proteins (HSPA1A, HSPA1B, HSPA6) and stress-responsive factors (DNAJB1, BAG3, IER5) are among the most highly upregulated. Downregulated genes (blue): Genes significantly decreased in expression (log2FC < -1.5, adjusted *P*-value < 0.05), including CCN1, SGK1, and AFP. Non-significant genes (black): Genes that do not meet statistical thresholds (|log2FC| ≤ 1.5 or adjusted *P*-value ≥ 0.05). The overall pattern suggests a strong heat shock response, with molecular chaperones and stress-associated genes strongly induced, while other pathways, potentially related to normal cellular function, are repressed.

#### Gene Ontology (GO) and KEGG Pathway Enrichment

Functional enrichment analysis using gene set enrichment analysis (GSEA) and topGO revealed key biological pathways involved in the heat shock response. Gene Ontology (GO) enrichment analysis identified a significant overrepresentation of cellular stress response pathways, including those related to molecular chaperone activity, unfolded protein response, and translational regulation (Figure 9, Supplementary Figure S5, and Supplementary Table S2).

**Figure 9.**
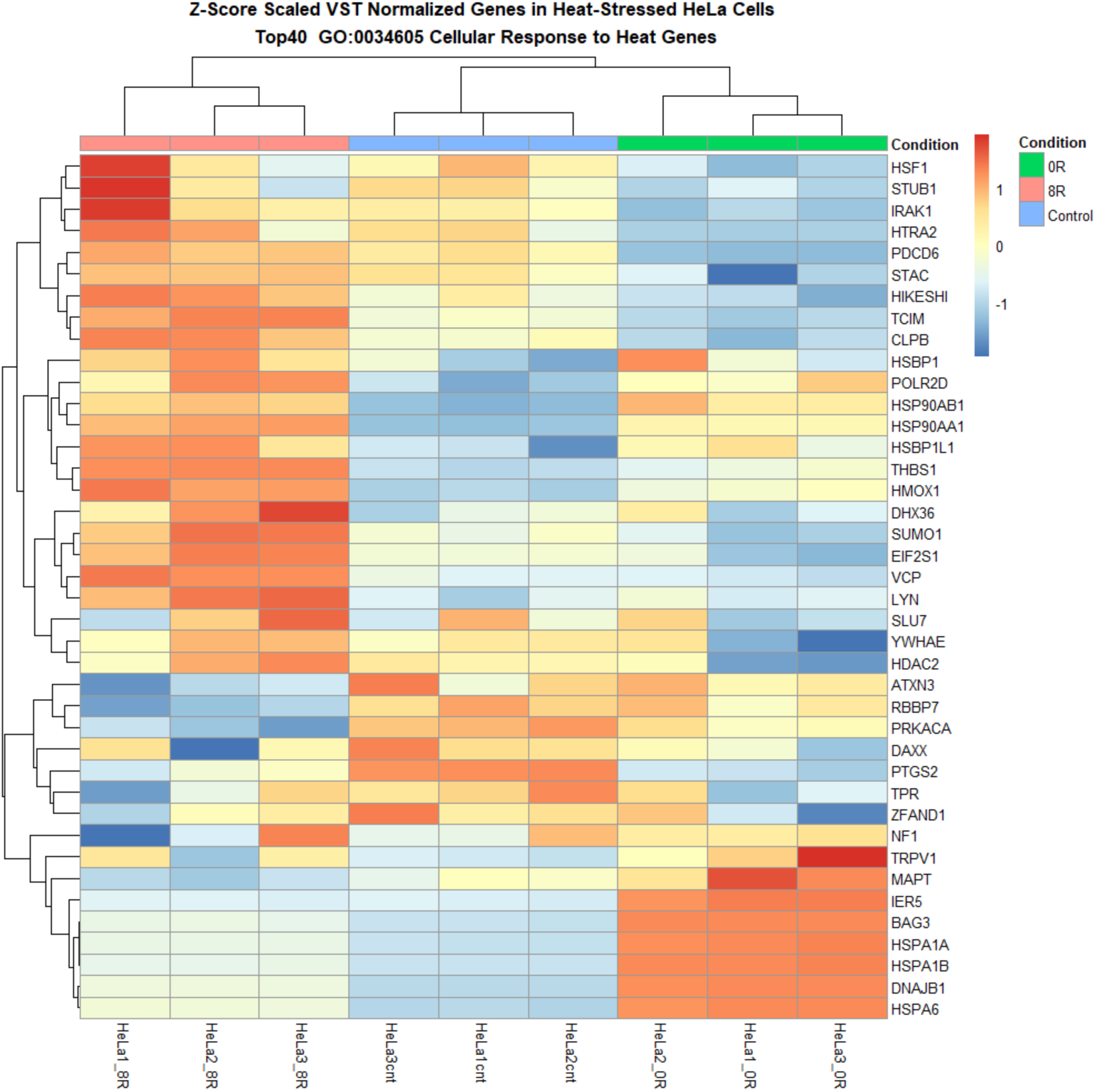
Heatmap of Differentially Expressed Heat Shock Response Genes reveals distinct transcriptional shifts. The heatmap illustrates the hierarchical clustering of differentially expressed genes (DEGs) across experimental conditions, with each column representing an individual sample and each row corresponding to a specific gene. Genes upregulated in response to heat shock are shown in red, while downregulated genes are displayed in blue. The clustering highlights distinct transcriptional shifts, with a pronounced separation between control and heat shock (R0) conditions and a partial return to baseline expression levels in the recovery phase (R8). Upregulated genes include key molecular chaperones, stress-response regulators, and transcription factors, reinforcing the activation of heat shock pathways. Downregulated genes primarily include those associated with normal cellular homeostasis, suggesting temporary suppression of non-essential functions under stress.

Additionally, KEGG pathway analysis highlighted ribosome biogenesis, sphingolipid metabolism, and oxidative stress response as enriched pathways in response to heat shock. The transcriptional signature of heat shock-exposed cells included genes associated with protein refolding and metabolic adaptation. Notably, pathways related to RNA processing, chromatin regulation, and ribosome function exhibited dynamic regulation across conditions, emphasizing the interplay between transcriptional and translational control in the heat shock response. However, no single pathway fully accounted for the observed lipidomic shifts, suggesting a complex regulatory network rather than a direct transcriptional control mechanism. These findings indicate that while heat shock broadly affects gene expression, lipid metabolism-specific transcriptional responses are more subtle, necessitating further integrative analyses.

### 2.4. Focus on Lipid Metabolism Genes

#### Targeted Enrichment Plots & Manual Verification of GO Analyses

Initial gene ontology (GO) and pathway enrichment analyses did not highlight lipid metabolism as a major transcriptional response to heat shock. This observation suggests that lipidomic changes may be regulated at levels beyond transcription. Therefore, we used a more targeted approach to investigate lipid-related pathways and genes that might have been overlooked in the broader analyses.

GO results were manually verified by extracting and analyzing genes associated with lipid-related biological processes, molecular functions, and cellular components. A custom script was used to reconstruct the human GO network locally and filter for lipid-related terms (Supplementary Table S3) [43]. This process identified 352 genes in R0 vs. control and 444 genes in R8 vs. R0 associated with lipid-related GO terms. Still, only a small subset met differential expression cutoffs (|log2FC| > 1.5, adjusted *P-*value < 0.05). These findings reinforce the idea that lipid metabolism genes are transcriptionally responsive but do not dominate the stress response signature.

#### Custom Gene Set Heatmap and LFC Analyses

A series of heatmaps and log fold change (LFC) analyses were performed on predefined lipid metabolism gene sets to refine our understanding further. Heatmaps of variance-stabilized transformed (VST) counts revealed that while most lipid metabolism-related genes exhibited relatively stable expression, a subset of 60 genes appeared to undergo differential expression in response to R0 (Figures 9 and 10 and Supplementary Figure S6). These genes included ACACA, FASN, PLA2G4C, LPIN1, and PCK1, which are known regulators of lipid biosynthesis, fatty acid metabolism, and phospholipid remodeling.

**Figure 10.**
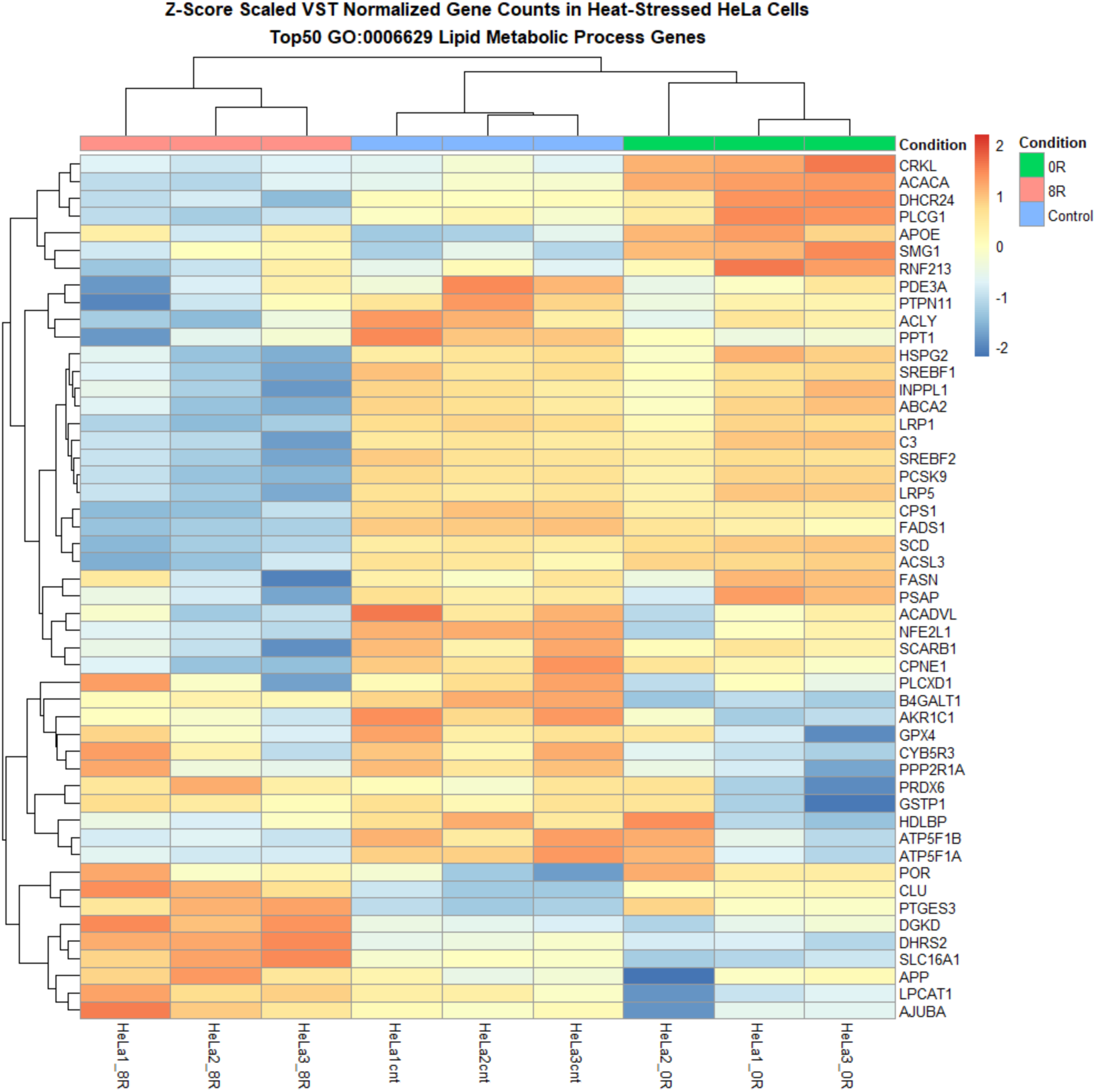
Heatmap of Lipid Metabolism-Related Genes highlights the hierarchical clustering of genes involved in lipid biosynthesis, remodeling, transport, and degradation. The heatmap presents the hierarchical clustering of lipid metabolism-related genes differentially expressed across experimental conditions. Each row corresponds to a gene associated with lipid biosynthesis, remodeling, transport, or degradation, while each column represents a sample. Genes upregulated in response to heat shock (R0) appear in red, whereas downregulated genes are displayed in blue.

However, quantitative evaluation using LFC values confirmed that only 13 genes met stringent differential expression cutoffs across multiple lipid metabolism pathways. These included SOCS1, CD74, ALOXE3, B3GALT4, and PLA2G4C (Supplementary Figure S6).

#### qPCR Validation of Lipid Metabolism Genes

Given the extensive lipidomic remodeling observed in response to heat shock, we performed qPCR (Figure 11) of key lipid metabolism-related genes to determine whether the transcriptional changes detected in RNA-seq correlated with lipid abundance shifts. The selection of genes was guided by specific lipidomic trends identified in our analysis. The observed increase in free fatty acids (FA) following heat shock suggested upregulation of *de novo* lipogenesis, warranting the inclusion of fatty acid synthase (FASN) in our validation panel. Additionally, the enrichment of unsaturated fatty acids indicated potential alterations in lipid desaturation pathways, leading us to examine fatty acid desaturase genes (FADS1 and FADS2).

**Figure 11.**
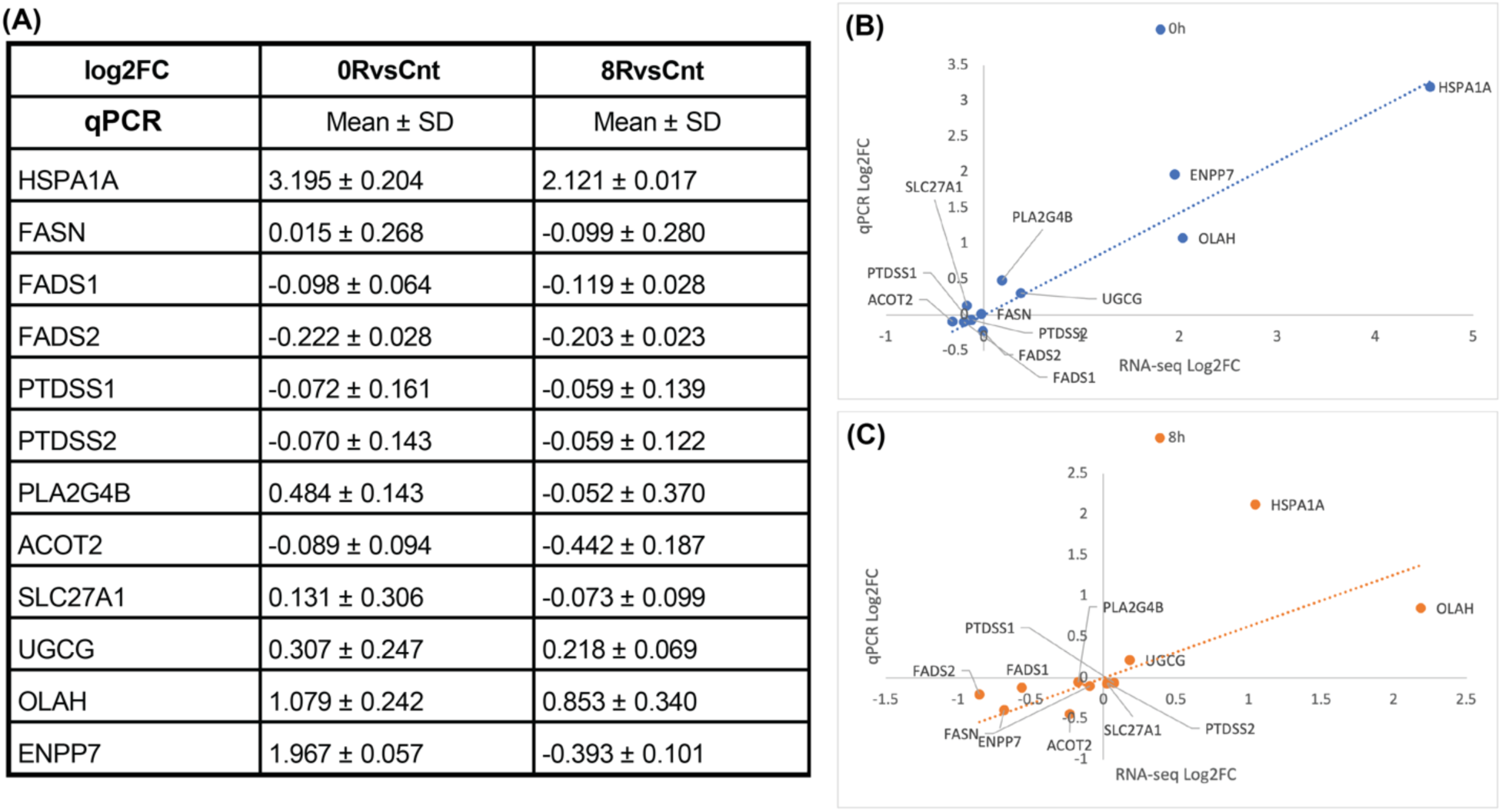
qPCR Validation of Lipid Metabolism Genes. (A) Quantitative panel summarizing the mean log₂ fold change (log₂FC) and standard deviation (SD) of selected lipid metabolism-related genes measured by qPCR for 0RvsCnt and 8RvsCnt comparisons. Most lipid-related genes exhibit minimal transcriptional changes, aligning with RNA-seq findings. (B, C) Scatter plots comparing log₂FC values obtained from RNA-seq and qPCR for the 0h (B, blue) and 8h (C, orange) conditions. A strong correlation is observed for HSPA1A, confirming heat shock-induced upregulation.

Similarly, since phosphatidylserine (PS) levels were significantly elevated in response to heat shock, we included phosphatidylserine synthase genes (PTDSS1 and PTDSS2) to determine whether their transcriptional regulation contributed to PS accumulation. Given the role of lipases in lipid remodeling, we analyzed phospholipase A2 group IVB (PLA2G4B), while genes associated with fatty acid oxidation (ACOT2, SLC27A1), sphingolipid biosynthesis (UGCG), and lipid hydrolysis (OLAH, ENPP7) were selected to assess broader metabolic changes.

Despite clear lipidomic shifts, qPCR results (Figure 11) confirmed that none of these genes exhibited substantial transcriptional changes, consistent with RNA-seq findings. This finding reinforces the notion that heat shock-induced lipid remodeling is not primarily driven by transcriptional regulation. Instead, post-transcriptional mechanisms, enzymatic activity modulation, and metabolic flux adjustments are likely responsible for the observed lipidomic phenotype.

These findings further support the need for multi-omics integration, where we examine whether joint lipidomic-transcriptomic analyses can reveal underlying regulatory mechanisms governing heat-induced lipid remodeling.

### 2.5. Multi-Omics Integration: Linking Lipidomics and Transcriptomics

#### Joint Pathway Analysis Reveals Coordinated Lipid-Gene Responses

To explore the interplay between lipidomic and transcriptomic responses to heat shock, we conducted a joint pathway analysis using MetaboAnalyst’s multi-omics integration module. This analysis incorporated differentially abundant lipids and differentially expressed genes (DEGs) across conditions to identify enriched metabolic pathways with concurrent transcriptional and lipidomic regulation.

The joint pathway analysis identified several key metabolic pathways significantly enriched in response to heat shock, including biosynthesis of unsaturated fatty acids, sphingolipid metabolism, and alpha-linolenic acid metabolism (Figure 12, Supplementary Figure S7, and Supplementary Table S4). Notably, glycerophospholipid metabolism was also highlighted, further supporting the role of membrane remodeling as a major adaptive mechanism to heat stress. These pathways were consistently enriched across multiple comparisons (R0 vs. control, R8 vs. control, and R8 vs. R0; Supplementary Table S4 and Supplementary Figure S7), suggesting a tightly regulated response coordinating lipid metabolism and gene expression.

**Figure 12.**
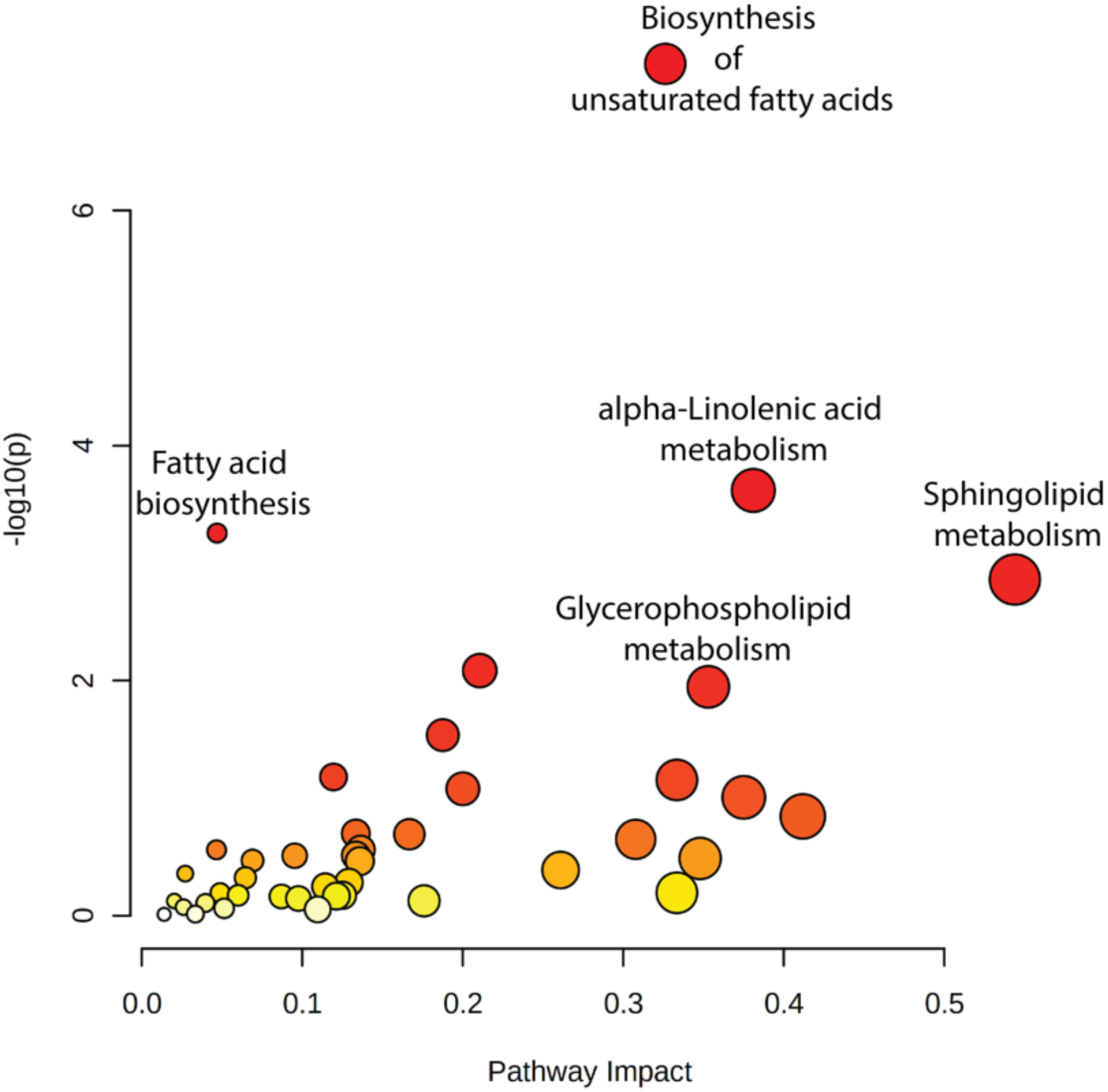
Integrated Pathway Analysis of Lipidomic and Transcriptomic Data identifies coordinated changes after heat shock. Key metabolic pathways, including biosynthesis of unsaturated fatty acids, alpha-linolenic acid metabolism, fatty acid biosynthesis, and sphingolipid metabolism, exhibit significant coordination between lipidomic and transcriptomic changes in response to heat shock. Dot size represents pathway enrichment, while color intensity corresponds to statistical significance (-log₁₀(*P-*value)), with darker red indicating higher significance.

#### Network Analysis Identifies Key Regulatory Nodes in Stress Adaptation

To further dissect interactions between lipid species and gene expression, we performed a network analysis to visualize molecular hubs integrating lipidomic and transcriptomic changes. This analysis identified key regulatory nodes in stress adaptation, highlighting genes and lipid species with high betweenness centrality in metabolic subnetworks (Figure 13).

**Figure 13.**
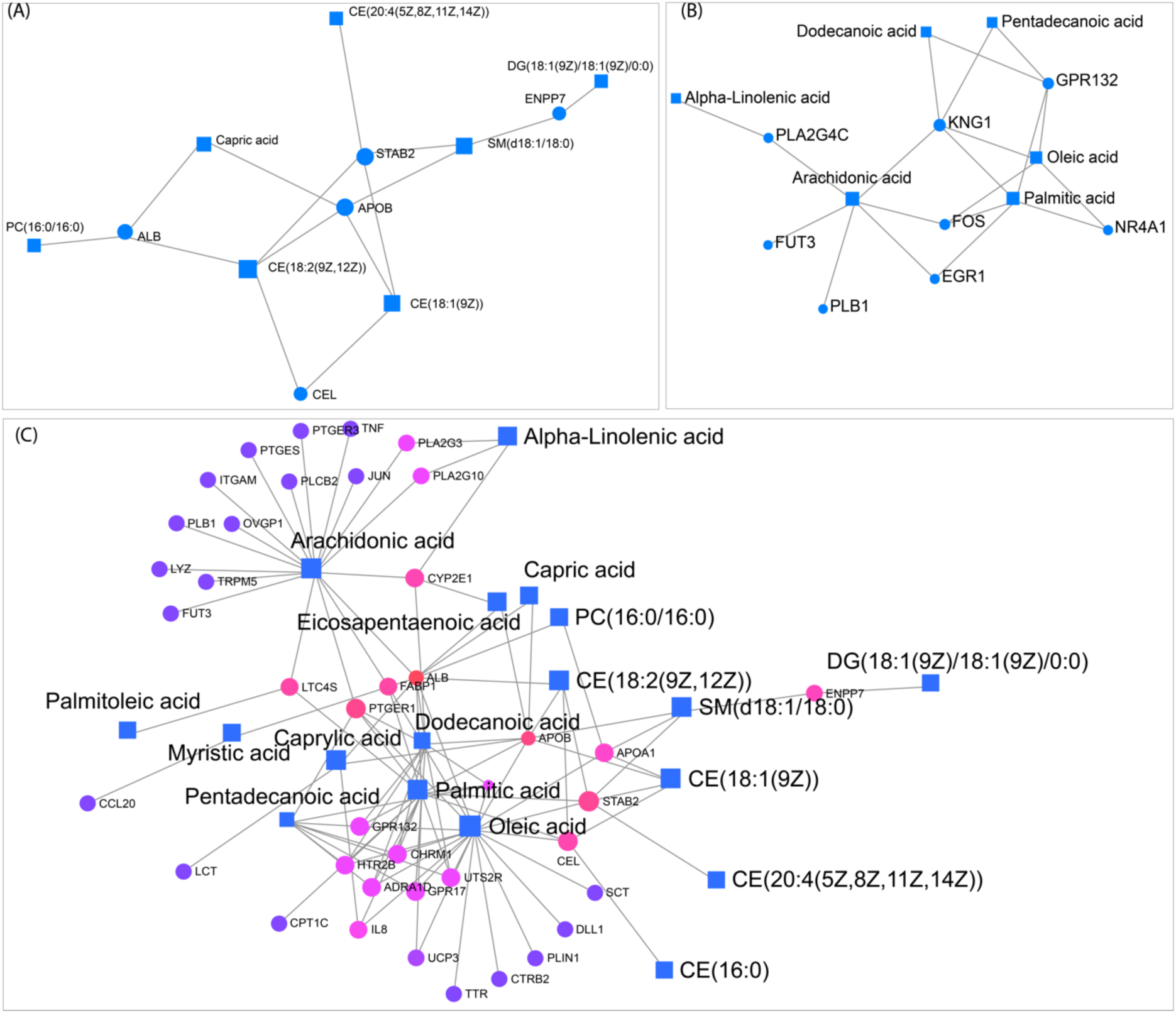
Lipid-Gene Interaction Network highlights key metabolic connections and transcriptional regulators associated with lipid metabolism. Network representations of lipidomic and transcriptomic interactions, where squares represent lipids and circles represent genes. (A) Displays lipid-protein interactions, highlighting associations between specific lipids such as PC(16:0/16:0) and capric acid with lipid-binding proteins, including ALB, APOB, and STAB2. (B) Shows metabolic connections of alpha-linolenic acid, arachidonic acid, and oleic acid with lipid-related genes such as PLA2G4C, GPR132, and NR4A1. (C) Expands the network to include additional lipids and genes, illustrating connections between fatty acids such as arachidonic acid, palmitic acid, and capric acid with multiple transcriptional regulators. Lines indicate interaction edges linking lipids and genes.

For the R0 vs. control comparison, network analysis revealed a core subnetwork centered around STAB2 and APOB, genes involved in lipid transport and metabolism (Figure 13A). These genes exhibited high betweenness centrality and were directly linked to sphingomyelin (SM) and cholesterol ester levels, which were significantly altered under heat shock (Figure 2 and Supplementary Figure S1). The enrichment of SM and cholesterol esters within this network suggests an adaptive response to maintain membrane integrity during acute stress. In the R8 vs control condition, KNG1 and arachidonic acid emerged as key regulatory nodes, linking lipid signaling pathways with inflammatory and stress-related responses (Figure 13B). Finally, in the R8 vs R0 network, ALB (albumin), APOB, STAB2, and PTGER1 formed central hubs, suggesting a role in lipid transport and metabolic recovery post-stress (Figure 13C). The network also highlighted arachidonic acid, oleic acid, palmitic acid, and other lipids as key metabolites with multiple gene interactions. These findings provide a systems-level perspective on how lipidome and transcriptome responses coordinate stress adaptation and membrane remodeling.

#### Overlapping Signatures Between Lipidomics and Transcriptomics

Despite the extensive lipid remodeling observed in response to heat shock, initial global transcriptomic enrichment analyses did not highlight lipid metabolism as a primary feature (Supplementary Figure S4).

The joint pathway and network analyses reinforce the importance of membrane lipid reorganization during heat shock adaptation. Although transcriptomic changes alone did not fully explain the lipidomic phenotype, multi-omics integration revealed functionally relevant pathways where gene expression changes align with lipidomic shifts (Figure 12 and Figure 13). These findings highlight the necessity of integrating molecular datasets to uncover regulatory mechanisms governing cellular adaptation to stress.

## 3. Discussion

The heat shock response (HSR) is a well-conserved adaptive mechanism that protects cells from proteotoxic stress by activating molecular chaperones, altering protein homeostasis, and triggering broad metabolic reprogramming [2,4,44]. While the transcriptional regulation of HSR is well characterized, the role of lipid remodeling in cellular stress adaptation remains underexplored. This study integrates lipidomics and transcriptomics to elucidate the metabolic adjustments accompanying heat shock, revealing that heat-induced lipidomic changes are not primarily driven by transcriptional regulation.

Lipidomics analysis revealed a global shift in lipid composition during heat shock, with specific enrichment of fatty acids, phospholipids, and sphingolipids. The significant increase in phosphatidylserine (PS) and phosphatidylethanolamine (PE) suggests a role for membrane restructuring, vesicle trafficking, and stress-induced lipid signaling [45]. Sphingolipid metabolism was also highly enriched, consistent with its known roles in cellular stress signaling, apoptosis, and membrane integrity regulation [46]. These lipid changes are likely to alter the biophysical properties of the plasma membrane (PM), such as fluidity and rigidity, which are essential for stress sensing and cellular adaptation [8,47-50]. Membrane composition is critical in modulating protein-lipid interactions and organizing signaling platforms. The sustained enrichment of PS and PE suggests prolonged alterations in membrane dynamics, which may affect protein localization, vesicle trafficking, and membrane-associated signaling events that facilitate stress adaptation [8,47-51].

Despite the significant lipidomic changes observed, pathway enrichment analyses did not identify a single dominant metabolic pathway driving the heat shock response. This finding suggests that lipid remodeling during heat stress is governed by dispersed metabolic adjustments rather than a single transcriptionally regulated program [21]. The lack of strong pathway enrichment may be due to the complexity of lipid metabolism, where multiple overlapping pathways contribute to lipidomic shifts [52], or due to limitations in annotation databases that do not fully capture the dynamic remodeling of lipids in acute stress conditions.

RNA-seq analysis identified thousands of differentially expressed genes (DEGs) responding to heat shock and recovery, clustering into distinct transcriptional response groups. As expected, genes involved in chaperone activity, unfolded protein response, and cellular stress signaling were among the most upregulated in R0, shifting toward metabolic adjustment during recovery (R8) [53,54]. However, global transcriptomic enrichment analyses did not highlight lipid metabolism as a dominant feature of the heat shock response. Despite the widespread lipidomic alterations observed, genes involved in fatty acid biosynthesis, phospholipid remodeling, and sphingolipid metabolism did not exhibit statistically significant transcriptional upregulation. The observation that lipid remodeling occurs without strong transcriptional enrichment suggests that additional layers of regulation—such as enzyme activity, metabolic flux adjustments, or post-transcriptional modifications—may be involved. Lipid metabolism is often controlled by substrate availability, enzyme activity, and post-translational modifications rather than transcriptional upregulation [18,19,21-26]. The sustained increase in fatty acids (FA), ceramides (Cer), phosphatidylserine (PS), phosphatidylethanolamine (PE), and sphingolipids in R8 further suggests that stress-induced lipid remodeling is not transient but persists beyond the immediate heat shock response, potentially through altered lipid trafficking and turnover.

Lipid remodeling is a key adaptive strategy across different biological systems to cope with thermal and metabolic stress. Previous studies have shown that heat stress induces a shift from membrane lipids to storage lipids as a protective mechanism against damage ([13,55]. In plants, lipid remodeling during heat stress is highly coordinated, adjusting membrane fluidity and metabolic fluxes to enhance survival [22,23,28-30,49]. Similarly, stress-induced lipidomic alterations in human cells impact membrane stability, protein trafficking, and energy metabolism [56]. The findings of this study align with these observations, reinforcing the idea that heat stress triggers lipidomic shifts essential for maintaining cellular integrity. The increase in phospholipids involved in membrane remodeling and sphingolipid-associated stress signaling suggests lipid metabolism is integral to cellular stress tolerance.

The metabolic rewiring observed in the HSR shares similarities with lipid metabolic adaptations in cancer. Cancer cells frequently reprogram lipid metabolism to support rapid proliferation, membrane biosynthesis, and survival under stress conditions [31]. For example, fatty acid metabolism and lipid storage mechanisms are upregulated in tumors to buffer oxidative and proteotoxic stress. Furthermore, cancer cells manipulate lipid metabolic pathways to evade cell death mechanisms such as ferroptosis, a form of iron-dependent lipid peroxidation [57]. The results suggest lipid remodeling during heat shock mirrors metabolic adjustments observed in cancer cells, particularly in phospholipid and sphingolipid metabolism. Understanding these lipidomic changes in a stress context may provide new insights into how cancer cells exploit lipid metabolism for survival and therapy resistance. Targeting lipid metabolic vulnerabilities in cancer—such as sphingolipid metabolism or fatty acid synthesis— could provide novel therapeutic strategies for disrupting stress-adaptive mechanisms in tumor cells.

Given the absence of potent lipid metabolic enrichment in global transcriptomic analyses, targeted enrichment analyses and manual verification of lipid-related pathways were performed. These analyses identified a subset of lipid metabolism genes with condition-dependent expression changes, including PCK1, PLA2G4C, and ACACA, which are involved in phospholipid biosynthesis and fatty acid metabolism. qPCR validation confirmed that these genes followed similar expression trends to RNA-seq, and the overall changes in expression were not large enough to account for the lipidomic phenotype. This finding reinforces the hypothesis that lipid remodeling during heat shock is not transcriptionally driven but rather controlled via post-transcriptional mechanisms, enzymatic activity, or metabolic flux adjustments.

Joint pathway and network analyses revealed that lipid remodeling is coordinated with stress adaptation pathways but does not strongly correlate with direct transcriptional changes. Integrating lipidomic and transcriptomic data identified key metabolic pathways with concurrent lipid and gene regulation, such as glycerophospholipid metabolism, sphingolipid biosynthesis, and fatty acid metabolism. While individual transcriptomic or lipidomic analyses did not fully explain the regulatory mechanisms underlying heat shock adaptation, the integration of both datasets revealed key metabolic pathways with concurrent lipidomic and transcriptional regulation. Network analysis identified central hubs linking lipid transport (APOB, STAB2), stress signaling (STAB2), and stress adaptation (KNG1, ALB), reinforcing the role of lipid remodeling in cellular resilience to thermal stress. These findings highlight the necessity of multi-omics approaches to uncover functionally relevant stress adaptation pathways that may not be apparent through single-layer analyses.

While this study provides new insights into lipidomic and transcriptomic integration in the heat shock response, some limitations should be acknowledged. A single transformed cell line (HeLa) may not fully represent lipidomic and transcriptomic responses in other cell types, mainly primary or non-cancerous cells. Additionally, while RNA-seq provided a broad overview of transcriptional changes, it does not capture post-transcriptional modifications or enzyme activity, which are critical regulators of lipid metabolism. The lipidomic analysis, while extensive, was constrained by database limitations, meaning some lipid species are not yet fully annotated, and consequently, not all measured lipids could be mapped to specific activity networks. Future studies can expand these findings to multiple cell types, integrate proteomics and metabolomics for a more complete picture of lipid regulation, and employ functional assays to validate key genes involved in lipid remodeling under heat stress.

Understanding how cells dynamically reprogram lipid metabolism in response to stress has broad implications beyond heat shock adaptation. Given the parallels between stress-induced lipid remodeling and cancer metabolism, further research into lipidomic regulation may uncover new therapeutic targets for diseases characterized by altered lipid homeostasis, including cancer and metabolic disorders.

## 4. Materials and Methods

### 4.1. Cell Culture

HeLa cells (ATCC® CCL-2™), originally derived from Henrietta Lacks, were obtained from ATCC in December 2016 and verified bi-annually. Cells were maintained in Minimum Essential Medium (MEM) supplemented with 10% fetal bovine serum (FBS), 2 mM L-glutamine, 0.1 mM non-essential amino acids (NEAA), 1 mM sodium pyruvate, and penicillin-streptomycin. Cultures were grown in a humidified atmosphere containing 5% CO₂ at 37°C and passaged every 2–3 days to maintain optimal growth.

### 4.2. Heat Shock Treatment

To examine lipidome changes following heat shock, HeLa cells were subjected to one of three conditions: control (37°C), heat shock (42°C for 60 minutes), or heat shock followed by recovery at 37°C for 8 hours. Cells were seeded into 175 cm² flasks and cultured to ∼80% confluency using low-passage cells (passages 4–7). For each experiment, three biological replicates per condition were prepared: one flask was maintained at 37°C as a control, while two flasks were subjected to heat shock. Of the heat-shocked flasks, one was harvested immediately after heat shock (0 hours recovery, 0R), and the other was allowed to recover at 37°C for 8 hours (8 hours recovery, 8R). Cells were harvested by trypsinization, washed with PBS, pelleted, and stored at -80°C for subsequent lipidomics and transcriptomics analyses.

### 4.3. Lipid Quantification and Analysis

#### Sample Preparation

Six biological replicates were prepared for lipidomic analysis, each containing four million HeLa cells. Cells were pelleted and stored at -80°C before use.

#### Mass Spectrometry and Lipidomics

Lipid extraction and analysis were conducted using a biphasic lipid extraction method at the UC Davis West Coast Metabolomics Center. Lipids were separated using a C18-based hybrid bridged column with a ternary water/acetonitrile/isopropanol gradient and analyzed on a ThermoFisher Scientific Q-Exactive HF mass spectrometer with electrospray ionization. MS/MS data were acquired in data-dependent mode, and accurate masses were normalized using constant reference ion infusion. Lipid annotations were processed using MS-DIAL vs. 4.90, with precursor mass errors <10 mDa, retention time matching, and MS/MS matching to corresponding lipid classes [58]. Lipid intensities were sum-normalized by representing each lipid as a fraction of total lipids within the sample, scaled by the median intensity for each treatment group [37]. Detailed results are provided in Supplementary Table S5.

#### Lipidomics Analysis

Mass spectrometry-identified lipids were annotated using PubchemPy and MetaboAnalyst to resolve identifiers from InChI Keys, SMILES, or common names into PubChem CID, HMDB, and KEGG formats. The provided InChI Keys were converted into PubChem identifiers using the "get_compounds" function in PubchemPy via custom scripts (https://github.com/lesolano/MS-Thesis). If automated resolution failed or discrepancies occurred, manual queries of PubChem, LipidMaps, and HMDB databases were performed to identify the best matches. MetaboAnalyst’s Metabolite ID Conversion tool further converted Pubchem identifiers to HMDB and KEGG formats for downstream analyses [59].

Lipid abundance data were visualized using boxplots and heatmaps to represent total and proportional lipid class abundances across experimental conditions. Heatmaps were generated with MetaboAnalyst’s hierarchical clustering tool, using Euclidean distances and Ward’s clustering method, to group samples by lipid profiles and highlight lipidomic changes. Boxplots depicted lipid abundance variations across major lipid classes [59].

Statistical analyses were conducted in MetaboAnalyst [59]. Univariate analyses included fold-change and Student’s t-tests to identify significantly altered lipids for each condition comparison (R0 vs. control, R8 vs. control, R8 vs. R0). Multivariate analyses involved one-way ANOVA with Fisher’s LSD post hoc tests to assess differences in lipid abundance across conditions, with results visualized as plots of lipid abundance vs. -log10(*P-*value). The false discovery rate (FDR)-adjusted *P-*value cutoff was set to 0.05 for all tests.

Dimensionality reduction techniques were applied to simplify the lipidomic dataset. Principal component analysis (PCA) was used to identify major sources of variation and estimate effect sizes, with results visualized as scree and 2D score plots. Hierarchical clustering heatmaps provided further insight into sample grouping and lipid group changes in response to heat shock.

Enrichment and pathway analyses were performed to explore metabolic pathway involvement. Qualitative over-representation analysis identified enriched metabolite sets and pathways using KEGG and SMPDB databases, producing bar charts, dot plots, and network views [59]. Quantitative pathway analyses further quantified pathway-level impacts, mapping lipid abundance to pathways and generating scatter plots and pathway network overlays. Analyses used KEGG and SMPDB libraries, with criteria for pathway inclusion set to at least two metabolites per pathway or set.

Final outputs from these analyses included detailed statistics, visualizations (e.g., bar charts, dot plots, scatter plots), and pathway maps that collectively highlighted lipidomic changes under heat shock and recovery conditions.

### 4.4. RNA Sequencing and Analysis

#### Sample Preparation, cDNA Library Preparation, and Sequencing

Cells from the control, 0R, and 8R conditions were sent to Novogene (Sacramento, CA) for RNA extraction, library preparation, and sequencing. Total RNA was isolated using the Qiagen RNeasy Kit. RNA quality and integrity were assessed using 1% agarose gel electrophoresis, a NanoPhotometer® spectrophotometer (IMPLEN, CA, USA), and the Agilent Bioanalyzer 2100 system. Sequencing libraries were constructed using the NEBNext® UltraTM RNA Library Prep Kit for Illumina® (NEB, USA).

Poly-A mRNA was isolated using oligo-dT magnetic beads, fragmented under elevated temperature, and reverse transcribed into cDNA. Second-strand cDNA synthesis used DNA Polymerase I and RNase H. The cDNA was end-repaired, adenylated, and ligated to NEBNext adaptors. Fragments of 150–200 bp were selected using AMPure XP beads (Beckman Coulter, USA) and amplified via PCR with Phusion High-Fidelity DNA polymerase and indexed primers. Library quality was validated on the Agilent Bioanalyzer 2100 system.

Paired-end sequencing was performed on the Illumina NovaSeq S4 PE100 platform, generating at least 30 million reads per sample. RNA-seq data were processed to ensure high-quality reads for differential gene expression analysis.

*Transcriptomics Analysis (additional detailed methods in [53])*

#### Read Processing and Quality Control

Paired-end sequencing reads in FASTQ.gz format were obtained from Novogene and processed on a Linux-based system using BBDuk for quality control. Reads were trimmed for adapters, filtered for PhiX sequences, and subjected to phred-based quality filtering (Q ≥ 10) to ensure high-quality inputs for downstream analyses [60,61]. Trimmed reads were saved with updated file naming conventions and accompanied by summary statistics for verification.

#### Reference Genome Indexing and Alignment

Reads were aligned to the human reference genome (GRCh38.p13) using the STAR aligner, which was configured for splice-aware two-pass mapping. Before alignment, the reference genome was indexed using Gencode primary assembly annotation files and STAR’s genome generation mode [62]. Alignment outputs included sorted BAM files, unmapped reads in FASTQ format, and gene count files for downstream analyses.

#### Gene Feature Counting

Feature counting was performed using HTSeq, which quantified mapped reads per gene from sorted and indexed BAM files. Input included the GRCh38.104 GTF annotation file, and outputs comprised raw gene count matrices annotated with gene IDs and expression values [63].

#### Differential Gene Expression Analysis

Raw gene count matrices were imported into R and analyzed with DESeq2. Gene expression levels were modeled using negative binomial regression to identify differentially expressed genes (DEGs) for three pairwise comparisons: R0 vs. control, R8 vs. control, and R8 vs. R0. Results included log2 fold change (LFC), *P-*values, and FDR-adjusted *P-*values, which were visualized as volcano plots using ggplot2 [64]. All data can be found in [53].

#### Dimensionality Reduction

Variance-stabilizing transformation (VST) was applied to normalize gene counts and mitigate heteroscedasticity. Principal component analysis (PCA) was used to identify primary sources of variance, visualized via 2D score plots, while heatmaps of Z-score-scaled VST data highlighted gene expression patterns and sample clustering [65].

#### Functional Enrichment and Gene Set Analysis

Competitive gene set enrichment analysis (GSEA) was performed using the Hallmark (H), curated (C2), computational (C4), ontology (C5), and oncogenic (C6) collections from MSigDB to identify enriched pathways. GSEA ranked genes by LFC and evaluated enrichment significance using normalized enrichment scores (NES). Outputs included dot plots of top pathways and enrichment statistics [66,67]. Gene Ontology (GO) analysis with topGO provided hierarchical insights into biological processes, molecular functions, and cellular components enriched in the data [66,67].

#### Custom Gene Set and Lipid-Specific Analyses

A curated list of lipid-related GO terms and pathways was generated by parsing GO and QuickGO databases. DESeq2-calculated LFC values and DEG statistics were mapped to these custom gene sets to identify transcriptional changes associated with lipid metabolism. Heatmaps of Z-score-scaled VST data were created for targeted lipid-related gene sets, highlighting key genes and pathways involved in lipid homeostasis during heat shock [65,68].

#### Summary and Visualization

All processing and analysis outputs, including summary statistics, gene set enrichment plots, heatmaps, PCA plots, and DEG results, were consolidated into Excel files using Power Query for streamlined comparisons across experimental conditions.

#### Quantitative Polymerase Chain Reaction (qPCR)

Following the manufacturer’s protocol, RNA was isolated using 4 million HeLa cells per condition (control cells, 0 h recovery, 8 h recovery; different batches from the ones used for RNA-seq) using the Direct-Zol RNA mini-prep Kit (ZymoResearch, Irvine, CA, USA). Following the manufacturer’s protocol, cDNA was synthesized from 1μg of total RNA using the Superscript IV First-Strand synthesis system (ThermoFisher Scientific, Waltham, MA, USA) and Oligo (dT)20 primers. cDNA samples were diluted to a concentration of 50 ng/μL. qPCR reactions were prepared with the Power SYBR™ Green PCR Master Mix (ThermoFisher Scientific, Waltham, MA, USA) according to the manufacturer’s instructions. Three biological replicates were run for each gene and condition [Gene names and primers (generated using NCBI’s primer-blast utility) are shown in Supplementary Table S6]. qPCR was performed using the CFX96 Touch Real-Time Detection System (Bio-Rad, Hercules, CA, United States). The relative normalized expression [69] of the raw transcript levels was calculated using the Livak method for each gene [70] using the software provided with the instrument [71]. The reference genes used in this method were ACTB and GAPDH. Statistical significance was assessed using one-way ANOVA (Analysis of Variance) followed by post-hoc Tukey HSD (Honestly Significant Difference) and Bonferroni tests. A *p* value < 0.05 was considered statistically significant. Results were plotted via boxplot using BoxPlotR [72].

### 4.5. Integration of Lipidomics and Transcriptomics

#### Joint Pathway Analysis

To uncover insights unavailable from single-dataset analyses, MetaboAnalyst’s joint pathway analysis module was used to integrate lipid abundance and gene expression data [59]. Inputs included quantitative lipid abundance data and differentially expressed genes (DEGs) with accompanying fold changes. Enrichment analysis utilized a hypergeometric test, while topology measurements were based on degree centrality. Combined gene and metabolite queries were cross-referenced with KEGG and SMPDB databases to identify enriched pathways [73-76]. Outputs included summary statistic files, enriched pathway plots, and network visualizations with genes and lipids highlighted. Pathways containing both gene and lipid data were prioritized for further analysis.

#### Network Analysis

Gene-metabolite interaction networks were constructed using MetaboAnalyst’s network analysis module [59]. This tool aggregated subnetworks of metabolites and genes based on known interaction networks. Subnetworks were defined as having at least three nodes and could be filtered by node type, regulation status (up/down), or database-specific queries (e.g., KEGG, Reactome, GO, motif). Outputs included pathway names, interaction counts, and *P-*values for filtered queries. The overall gene-metabolite interaction network was visualized and saved as an image for interpretation.

## Supplementary Materials

The manuscript includes seven supplementary figures (S1-S7) and five supplementary tables (S1-S6).

## Author Contributions

Conceptualization, NN and LS; Data curation, LS and NN; Formal analysis, LS, AR, YC, WDH, and NN; Funding acquisition, NN; Methodology, LS and NN; Resources, NN; Writing – original draft, LS and NN; Writing – review & editing, LS, AR, YC, WDH, and NN.

## Data Availability Statement

All data reported are provided in the text and supplemental materials. The raw sequencing data are hosted at NCBI (GEO: GSE285497). The scripts used can be found at (https://github.com/lesolano/MS-Thesis).

## Acknowledgments

We thank Dr. Dimitra Chalkia for her valuable comments and help with the analysis of the manuscript.

## References Cited

1. Pessa, J.C.; Joutsen, J.; Sistonen, L. Transcriptional reprogramming at the intersection of the heat shock response and proteostasis. Mol Cell 2024, 84, 80–93, doi:10.1016/j.molcel.2023.11.024.

2. Richter, K.; Haslbeck, M.; Buchner, J. The heat shock response: life on the verge of death. Mol Cell 2010, 40, 253–266, doi:S1097-2765(10)00782-3 [pii] 10.1016/j.molcel.2010.10.006.

3. Morimoto, R.I. Regulation of the heat shock transcriptional response: cross talk between a family of heat shock factors, molecular chaperones, and negative regulators. Genes Dev 1998, 12, 3788–3796, doi:10.1101/gad.12.24.3788.

4. Bukau, B.; Weissman, J.; Horwich, A. Molecular chaperones and protein quality control. Cell 2006, 125, 443–451, doi:10.1016/j.cell.2006.04.014.

5. Horvath, I.; Glatz, A.; Nakamoto, H.; Mishkind, M.L.; Munnik, T.; Saidi, Y.; Goloubinoff, P.; Harwood, J.L.; Vigh, L. Heat shock response in photosynthetic organisms: membrane and lipid connections. Prog Lipid Res 2012, 51, 208–220, doi:10.1016/j.plipres.2012.02.002.

6. Johnston, M.K.; Jacob, N.P.; Brodl, M.R. Heat shock-induced changes in lipid and protein metabolism in the endoplasmic reticulum of barley aleurone layers. Plant Cell Physiol 2007, 48, 31–41, doi:10.1093/pcp/pcl037.

7. Narayanan, S.; Zoong-Lwe, Z.S.; Gandhi, N.; Welti, R.; Fallen, B.; Smith, J.R.; Rustgi, S. Comparative Lipidomic Analysis Reveals Heat Stress Responses of Two Soybean Genotypes Differing in Temperature Sensitivity. Plants (Basel) 2020, 9, doi:10.3390/plants9040457.

8. Török, Z.; Crul, T.; Maresca, B.; Schütz, G.J.; Viana, F.; Dindia, L.; Piotto, S.; Brameshuber, M.; Balogh, G.; Péter, M.;, et al. Plasma membranes as heat stress sensors: from lipid-controlled molecular switches to therapeutic applications. Biochim Biophys Acta 2014, 1838, 1594–1618, doi:10.1016/j.bbamem.2013.12.015.

9. Chauve, L.; Hodge, F.; Murdoch, S.; Masoudzadeh, F.; Mann, H.J.; Lopez-Clavijo, A.F.; Okkenhaug, H.; West, G.; Sousa, B.C.; Segonds-Pichon, A.;, et al. Neuronal HSF-1 coordinates the propagation of fat desaturation across tissues to enable adaptation to high temperatures in C. elegans. PLoS Biol 2021, 19, e3001431, doi:10.1371/journal.pbio.3001431.

10. Kuan, Y.C.; Hashidume, T.; Shibata, T.; Uchida, K.; Shimizu, M.; Inoue, J.; Sato, R. Heat Shock Protein 90 Modulates Lipid Homeostasis by Regulating the Stability and Function of Sterol Regulatory Element-binding Protein (SREBP) and SREBP Cleavage-activating Protein. J Biol Chem 2017, 292, 3016–3028, doi:10.1074/jbc.M116.767277.

11. Harayama, T.; Riezman, H. Understanding the diversity of membrane lipid composition. Nat Rev Mol Cell Biol 2018, 19, 281–296, doi:10.1038/nrm.2017.138.

12. van Meer, G.; Voelker, D.R.; Feigenson, G.W. Membrane lipids: where they are and how they behave. Nature Reviews Molecular Cell Biology 2008, 9, 112–124, doi:10.1038/nrm2330.

13. Jarc, E.; Petan, T. Lipid Droplets and the Management of Cellular Stress. Yale J Biol Med 2019, 92, 435–452.

14. Kim, H.S.; Kim, M.; Park, W.K.; Chang, Y.K. Enhanced Lipid Production of Chlorella sp. HS2 Using Serial Optimization and Heat Shock. J Microbiol Biotechnol 2020, 30, 136-145, doi:10.4014/jmb.1910.10033.

15. Klose, C.; Surma, M.A.; Gerl, M.J.; Meyenhofer, F.; Shevchenko, A.; Simons, K. Flexibility of a eukaryotic lipidome--insights from yeast lipidomics. PLoS One 2012, 7, e35063, doi:10.1371/journal.pone.0035063.

16. Lingwood, D.; Simons, K. Lipid rafts as a membrane-organizing principle. Science 2010, 327, 46–50, doi:10.1126/science.1174621.

17. Peksel, B.; Gombos, I.; Peter, M.; Vigh, L., Jr.; Tiszlavicz, A.; Brameshuber, M.; Balogh, G.; Schutz, G.J.; Horvath, I.; Vigh, L.;, et al. Mild heat induces a distinct "eustress" response in Chinese Hamster Ovary cells but does not induce heat shock protein synthesis. Sci Rep 2017, 7, 15643, doi:10.1038/s41598-017-15821-8.

18. Bromberg, Z.; Weiss, Y. The Role of the Membrane-Initiated Heat Shock Response in Cancer. Front Mol Biosci 2016, 3, 12, doi:10.3389/fmolb.2016.00012.

19. Sun, A.Z.; Chen, L.S.; Tang, M.; Chen, J.H.; Li, H.; Jin, X.Q.; Yi, Y.; Guo, F.Q. Lipidomic Remodeling in Begonia grandis Under Heat Stress. Front Plant Sci 2022, 13, 843942, doi:10.3389/fpls.2022.843942.

20. Jenkins, G.M.; Richards, A.; Wahl, T.; Mao, C.; Obeid, L.; Hannun, Y. Involvement of yeast sphingolipids in the heat stress response of Saccharomyces cerevisiae. J Biol Chem 1997, 272, 32566–32572, doi:10.1074/jbc.272.51.32566.

21. Krawczyk, H.E.; Rotsch, A.H.; Herrfurth, C.; Scholz, P.; Shomroni, O.; Salinas-Riester, G.; Feussner, I.; Ischebeck, T. Heat stress leads to rapid lipid remodeling and transcriptional adaptations in Nicotiana tabacum pollen tubes. Plant Physiol 2022, 189, 490–515, doi:10.1093/plphys/kiac127.

22. Amjadi, Z.; Hamzehzarghani, H.; Rodriguez, V.M.; Huang, Y.J.; Farahbakhsh, F. Studying temperature’s impact on Brassica napus resistance to identify key regulatory mechanisms using comparative metabolomics. Sci Rep 2024, 14, 19865, doi:10.1038/s41598-024-68345-3.

23. Higashi, Y.; Okazaki, Y.; Myouga, F.; Shinozaki, K.; Saito, K. Landscape of the lipidome and transcriptome under heat stress in Arabidopsis thaliana. Sci Rep 2015, 5, 10533, doi:10.1038/srep10533.

24. Estevao, I.L.; Kazman, J.B.; Bramer, L.M.; Nicora, C.; Ren, M.Q.; Sambuughin, N.; Munoz, N.; Kim, Y.M.; Bloodsworth, K.; Richert, M.;, et al. The impact of heat stress on the human plasma lipidome. Res Sq 2024, doi:10.21203/rs.3.rs-4548154/v1.

25. Estevao, I.L.; Kazman, J.B.; Bramer, L.M.; Nicora, C.; Ren, M.Q.; Sambuughin, N.; Munoz, N.; Kim, Y.M.; Bloodsworth, K.; Richert, M.;, et al. The human plasma lipidome response to exertional heat tolerance testing. Lipids Health Dis 2024, 23, 380, doi:10.1186/s12944-024-02322-7.

26. Zhang, J.; Fan, N.; Peng, Y. Heat shock protein 70 promotes lipogenesis in HepG2 cells. Lipids Health Dis 2018, 17, 73, doi:10.1186/s12944-018-0722-8.

27. Bach, L.; Faure, J.D. Role of very-long-chain fatty acids in plant development, when chain length does matter. C R Biol 2010, 333, 361–370, doi:10.1016/j.crvi.2010.01.014.

28. Barrero-Sicilia, C.; Silvestre, S.; Haslam, R.P.; Michaelson, L.V. Lipid remodelling: Unravelling the response to cold stress in Arabidopsis and its extremophile relative Eutrema salsugineum. Plant Sci 2017, 263, 194–200, doi:10.1016/j.plantsci.2017.07.017.

29. Spivey, W.W.; Rustgi, S.; Welti, R.; Roth, M.R.; Burow, M.D.; Bridges, W.C.; Narayanan, S. Lipid modulation contributes to heat stress adaptation in peanut. Frontiers in Plant Science 2023, 14, doi:10.3389/fpls.2023.1299371.

30. Cantarero, S.I.; Flores, E.; Allbrook, H.; Aguayo, P.; Vargas, C.A.; Tamanaha, J.E.; Scholz, J.B.C.; Bach, L.T.; Löscher, C.R.; Riebesell, U.;, et al. Lipid remodeling in phytoplankton exposed to multi-environmental drivers in a mesocosm experiment. Biogeosciences 2024, 21, 3927–3958, doi:10.5194/bg-21-3927-2024.

31. Beloribi-Djefaflia, S.; Vasseur, S.; Guillaumond, F. Lipid metabolic reprogramming in cancer cells. Oncogenesis 2016, 5, e189–e189, doi:10.1038/oncsis.2015.49.

32. Baenke, F.; Peck, B.; Miess, H.; Schulze, A. Hooked on fat: the role of lipid synthesis in cancer metabolism and tumour development. Dis Model Mech 2013, 6, 1353–1363, doi:10.1242/dmm.011338.

33. Koundouros, N.; Poulogiannis, G. Reprogramming of fatty acid metabolism in cancer. Br J Cancer 2020, 122, 4–22, doi:10.1038/s41416-019-0650-z.

34. Munir, R.; Lisec, J.; Swinnen, J.V.; Zaidi, N. Lipid metabolism in cancer cells under metabolic stress. British Journal of Cancer 2019, 120, 1090–1098, doi:10.1038/s41416-019-0451-4.

35. Wang, W.; Bai, L.; Li, W.; Cui, J. The Lipid Metabolic Landscape of Cancers and New Therapeutic Perspectives. Frontiers in Oncology 2020, 10, doi:10.3389/fonc.2020.605154.

36. van der Veen, J.N.; Kennelly, J.P.; Wan, S.; Vance, J.E.; Vance, D.E.; Jacobs, R.L. The critical role of phosphatidylcholine and phosphatidylethanolamine metabolism in health and disease. Biochimica et Biophysica Acta (BBA) - Biomembranes 2017, 1859, 1558–1572, 10.1016/j.bbamem.2017.04.006.

37. Garcia, G.; Zhang, H.; Moreno, S.; Tsui, C.K.; Webster, B.M.; Higuchi-Sanabria, R.; Dillin, A. Lipid homeostasis is essential for a maximal ER stress response. Elife 2023, 12, doi:10.7554/eLife.83884.

38. Sah, S.K.; Sofo, A. Editorial: The role of lipids in abiotic stress responses. Frontiers in Plant Science 2024, 15, doi:10.3389/fpls.2024.1378485.

39. Legeret, B.; Schulz-Raffelt, M.; Nguyen, H.M.; Auroy, P.; Beisson, F.; Peltier, G.; Blanc, G.; Li-Beisson, Y. Lipidomic and transcriptomic analyses of Chlamydomonas reinhardtii under heat stress unveil a direct route for the conversion of membrane lipids into storage lipids. Plant Cell Environ 2016, 39, 834–847, doi:10.1111/pce.12656.

40. Reich, S.; Nguyen, C.D.L.; Has, C.; Steltgens, S.; Soni, H.; Coman, C.; Freyberg, M.; Bichler, A.; Seifert, N.; Conrad, D.;, et al. A multi-omics analysis reveals the unfolded protein response regulon and stress-induced resistance to folate-based antimetabolites. Nature Communications 2020, 11, 2936, doi:10.1038/s41467-020-16747-y.

41. Mengelkoch, S.; Gassen, J.; Lev-Ari, S.; Alley, J.C.; Schüssler-Fiorenza Rose, S.M.; Snyder, M.P.; Slavich, G.M. Multi-omics in stress and health research: study designs that will drive the field forward. Stress 2024, 27, 2321610, doi:10.1080/10253890.2024.2321610.

42. Chaurasia, B.; Summers, S.A. Ceramides in Metabolism: Key Lipotoxic Players. Annu Rev Physiol 2021, 83, 303–330, doi:10.1146/annurev-physiol-031620-093815.

43. Solano, L.E.; D’Sa, N.M.; Nikolaidis, N. PRRGO: A Tool for Visualizing and Mapping Globally Expressed Genes in Public Gene Expression Omnibus RNA-Sequencing Studies to PageRank-scored Gene Ontology Terms. bioRxiv 2024, doi:10.1101/2024.01.21.576540.

44. Horowitz, M.; Robinson, S.D. Heat shock proteins and the heat shock response during hyperthermia and its modulation by altered physiological conditions. Prog Brain Res 2007, 162, 433–446.

45. Horn, A.; Jaiswal, J.K. Structural and signaling role of lipids in plasma membrane repair. Curr Top Membr 2019, 84, 67–98, doi:10.1016/bs.ctm.2019.07.001.

46. Ogretmen, B. Sphingolipid metabolism in cancer signalling and therapy. Nat Rev Cancer 2018, 18, 33–50, doi:10.1038/nrc.2017.96.

47. Wu, G.; Baumeister, R.; Heimbucher, T. Molecular Mechanisms of Lipid-Based Metabolic Adaptation Strategies in Response to Cold. Cells 2023, 12, doi:10.3390/cells12101353.

48. Szlasa, W.; Zendran, I.; Zalesińska, A.; Tarek, M.; Kulbacka, J. Lipid composition of the cancer cell membrane. Journal of Bioenergetics and Biomembranes 2020, 52, 321–342, doi:10.1007/s10863-020-09846-4.

49. Cano-Ramirez, D.L.; Carmona-Salazar, L.; Morales-Cedillo, F.; Ramírez-Salcedo, J.; Cahoon, E.B.; Gavilanes-Ruíz, M. Plasma Membrane Fluidity: An Environment Thermal Detector in Plants. Cells 2021, 10, doi:10.3390/cells10102778.

50. Zhang, T.; Hu, W.; Chen, W. Plasma Membrane Integrates Biophysical and Biochemical Regulation to Trigger Immune Receptor Functions. Front Immunol 2021, 12, 613185, doi:10.3389/fimmu.2021.613185.

51. Low, J.; Altman, R.; Badolian, A.; Cuaresma, A.B.; Briseño, C.; Keshet, U.; Fiehn, O.; Stahelin, R.V.; Nikolaidis, N. Heat-Induced Phosphatidylserine Changes Drive HSPA1A’s Plasma Membrane Localization. bioRxiv 2024, 2024.2012.2002.626454, doi:10.1101/2024.12.02.626454.

52. Alves, M.A.; Lamichhane, S.; Dickens, A.; McGlinchey, A.; Ribeiro, H.C.; Sen, P.; Wei, F.; Hyötyläinen, T.; Orešič, M. Systems biology approaches to study lipidomes in health and disease. Biochimica et Biophysica Acta (BBA) - Molecular and Cell Biology of Lipids 2021, 1866, 158857, 10.1016/j.bbalip.2020.158857.

53. Reinschmidt, A.; Solano, L.; Chavez, Y.; Hulsy, W.D.; Nikolaidis, N. Transcriptomics Unveil Canonical and Non-Canonical Heat Shock-Induced Pathways in Human Cell Lines. Int. J. Mol. Sci. 2025, 26, 1057.

54. Alagar Boopathy, L.R.; Jacob-Tomas, S.; Alecki, C.; Vera, M. Mechanisms tailoring the expression of heat shock proteins to proteostasis challenges. J Biol Chem 2022, 298, 101796, doi:10.1016/j.jbc.2022.101796.

55. Raetz, C.R.H.; Guan, Z.; Ingram, B.O.; Six, D.A.; Song, F.; Wang, X.; Zhao, J. Discovery of new biosynthetic pathways: the lipid A story. Journal of Lipid Research 2009, 50, S103–S108, doi:10.1194/jlr.R800060-JLR200.

56. Levental, K.R.; Malmberg, E.; Symons, J.L.; Fan, Y.-Y.; Chapkin, R.S.; Ernst, R.; Levental, I. Lipidomic and biophysical homeostasis of mammalian membranes counteracts dietary lipid perturbations to maintain cellular fitness. Nature Communications 2020, 11, 1339, doi:10.1038/s41467-020-15203-1.

57. Stockwell, B.R.; Friedmann Angeli, J.P.; Bayir, H.; Bush, A.I.; Conrad, M.; Dixon, S.J.; Fulda, S.; Gascón, S.; Hatzios, S.K.; Kagan, V.E.;, et al. Ferroptosis: A Regulated Cell Death Nexus Linking Metabolism, Redox Biology, and Disease. Cell 2017, 171, 273–285, doi:10.1016/j.cell.2017.09.021.

58. Cajka, T.; Fiehn, O. Increasing lipidomic coverage by selecting optimal mobile-phase modifiers in LC–MS of blood plasma. Metabolomics 2016, 12, 34, doi:10.1007/s11306-015-0929-x.

59. Chong, J.; Wishart, D.S.; Xia, J. Using MetaboAnalyst 4.0 for Comprehensive and Integrative Metabolomics Data Analysis. Curr Protoc Bioinformatics 2019, 68, e86, doi:10.1002/cpbi.86.

60. Grigoriev, I.V.; Nordberg, H.; Shabalov, I.; Aerts, A.; Cantor, M.; Goodstein, D.; Kuo, A.; Minovitsky, S.; Nikitin, R.; Ohm, R.A.;, et al. The genome portal of the Department of Energy Joint Genome Institute. Nucleic Acids Res 2012, 40, D26–32, doi:10.1093/nar/gkr947.

61. Nordberg, H.; Cantor, M.; Dusheyko, S.; Hua, S.; Poliakov, A.; Shabalov, I.; Smirnova, T.; Grigoriev, I.V.; Dubchak, I. The genome portal of the Department of Energy Joint Genome Institute: 2014 updates. Nucleic Acids Res 2014, 42, D26–31, doi:10.1093/nar/gkt1069.

62. Dobin, A.; Davis, C.A.; Schlesinger, F.; Drenkow, J.; Zaleski, C.; Jha, S.; Batut, P.; Chaisson, M.; Gingeras, T.R. STAR: ultrafast universal RNA-seq aligner. Bioinformatics 2013, 29, 15–21, doi:10.1093/bioinformatics/bts635.

63. Anders, S.; Pyl, P.T.; Huber, W. HTSeq--a Python framework to work with high-throughput sequencing data. Bioinformatics 2015, 31, 166–169, doi:10.1093/bioinformatics/btu638.

64. Ewels, P.; Magnusson, M.; Lundin, S.; Käller, M. MultiQC: summarize analysis results for multiple tools and samples in a single report. Bioinformatics 2016, 32, 3047–3048, doi:10.1093/bioinformatics/btw354.

65. Love, M.I.; Huber, W.; Anders, S. Moderated estimation of fold change and dispersion for RNA-seq data with DESeq2. Genome Biol 2014, 15, 550, doi:10.1186/s13059-014-0550-8.

66. Liberzon, A.; Subramanian, A.; Pinchback, R.; Thorvaldsdóttir, H.; Tamayo, P.; Mesirov, J.P. Molecular signatures database (MSigDB) 3.0. Bioinformatics 2011, 27, 1739–1740, doi:10.1093/bioinformatics/btr260.

67. Subramanian, A.; Tamayo, P.; Mootha, V.K.; Mukherjee, S.; Ebert, B.L.; Gillette, M.A.; Paulovich, A.; Pomeroy, S.L.; Golub, T.R.; Lander, E.S.;, et al. Gene set enrichment analysis: a knowledge-based approach for interpreting genome-wide expression profiles. Proc Natl Acad Sci U S A 2005, 102, 15545–15550, doi:10.1073/pnas.0506580102.

68. Durinck, S.; Spellman, P.T.; Birney, E.; Huber, W. Mapping identifiers for the integration of genomic datasets with the R/Bioconductor package biomaRt. Nat Protoc 2009, 4, 1184-1191, doi:10.1038/nprot.2009.97.

69. Vandesompele, J.; De Preter, K.; Pattyn, F.; Poppe, B.; Van Roy, N.; De Paepe, A.; Speleman, F. Accurate normalization of real-time quantitative RT-PCR data by geometric averaging of multiple internal control genes. Genome Biol 2002, 3, RESEARCH0034, doi:10.1186/gb-2002-3-7-research0034.

70. Livak, K.J.; Schmittgen, T.D. Analysis of relative gene expression data using real-time quantitative PCR and the 2(-Delta Delta C(T)) Method. Methods 2001, 25, 402–408, doi:10.1006/meth.2001.1262.

71. Hellemans, J.; Mortier, G.; De Paepe, A.; Speleman, F.; Vandesompele, J. qBase relative quantification framework and software for management and automated analysis of real-time quantitative PCR data. Genome Biol 2007, 8, R19, doi:10.1186/gb-2007-8-2-r19.

72. Spitzer, M.; Wildenhain, J.; Rappsilber, J.; Tyers, M. BoxPlotR: a web tool for generation of box plots. Nat Methods 2014, 11, 121–122, doi:10.1038/nmeth.2811.

73. Jewison, T.; Su, Y.; Disfany, F.M.; Liang, Y.; Knox, C.; Maciejewski, A.; Poelzer, J.; Huynh, J.; Zhou, Y.; Arndt, D.;, et al. SMPDB 2.0: big improvements to the Small Molecule Pathway Database. Nucleic Acids Res 2014, 42, D478–484, doi:10.1093/nar/gkt1067.

74. Kanehisa, M. Toward understanding the origin and evolution of cellular organisms. Protein Sci 2019, 28, 1947–1951, doi:10.1002/pro.3715.

75. Kanehisa, M.; Furumichi, M.; Sato, Y.; Kawashima, M.; Ishiguro-Watanabe, M. KEGG for taxonomy-based analysis of pathways and genomes. Nucleic Acids Res 2022, doi:10.1093/nar/gkac963.

76. Kanehisa, M.; Goto, S. KEGG: kyoto encyclopedia of genes and genomes. Nucleic Acids Res 2000, 28, 27–30, doi:10.1093/nar/28.1.27.

