## supplementary figures for "Dynamic Lipidome Reorganization in Response to Heat Shock Stress"

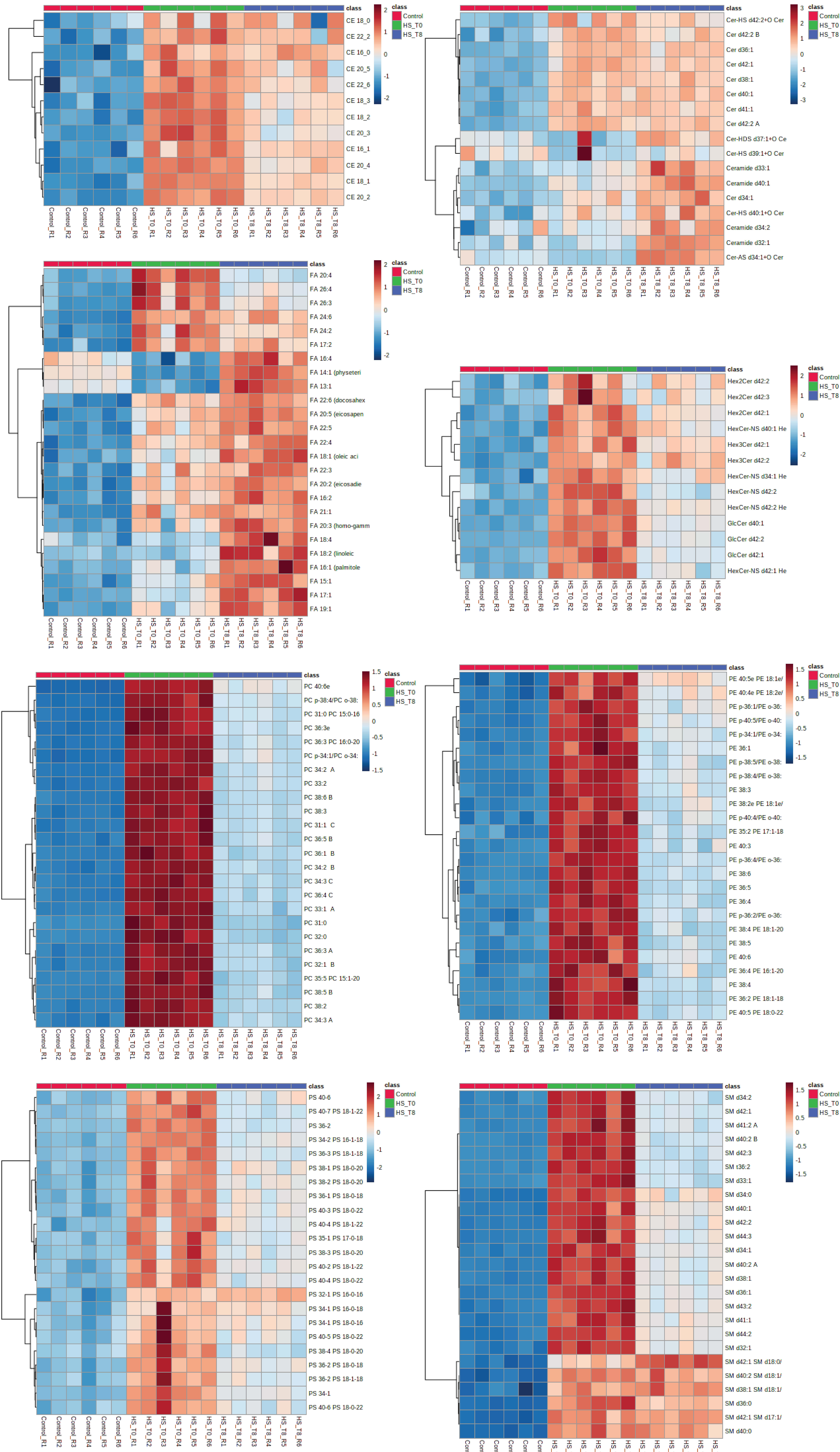


**Supplementary Figure S1**

**Supplementary Figure S1.** Global Heatmap of Lipidomic Changes. Expanded heatmaps displaying representative lipid species, organized by hierarchical clustering. Each heatmap represents a specific lipid class, showing relative abundance across experimental conditions (Control, HS_T0, and HS_T8). Lipid classes include cholesterol esters (CE), ceramides (Cer), fatty acids (FA; selected), hexosylceramides and Glyceroceramides (HexCer, GlcCer), phosphatidylcholines (PC; top 25), phosphatidylethanolamines (PE; top 25), phosphatidylserines (PS), and sphingomyelins (SM). Red indicates increased abundance, while blue represents decreased abundance. Distinct clustering patterns reveal lipid subclasses with transient or sustained changes in response to heat shock and recovery.

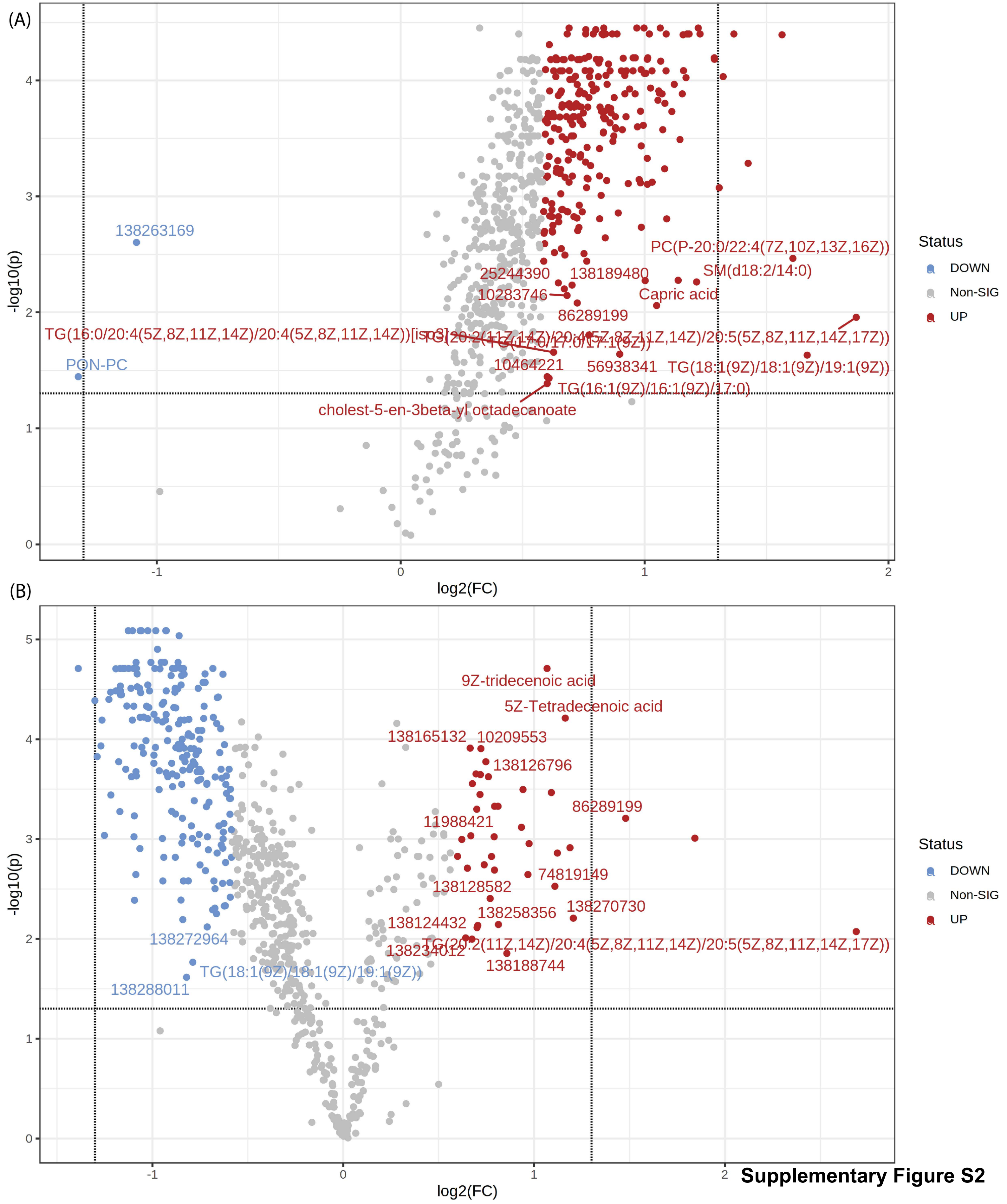


**Supplementary Figure S2**

**Supplementary Figure S2.** Differential Lipid Abundance in Recovery vs. Control. Volcano plots displaying lipid abundance changes post-recovery. (A) R8 vs. Control, highlighting significantly upregulated (red) and downregulated (blue) lipids, with non-significant lipids in gray. (B) R8 vs. R0, illustrating the extent of lipidomic shifts from heat shock to recovery. Dotted lines indicate significance thresholds for fold change ($|log2FC| > 1$) and adjusted p-value ($< 0.05$). PubChem CID identifiers label select lipids with the most pronounced changes.

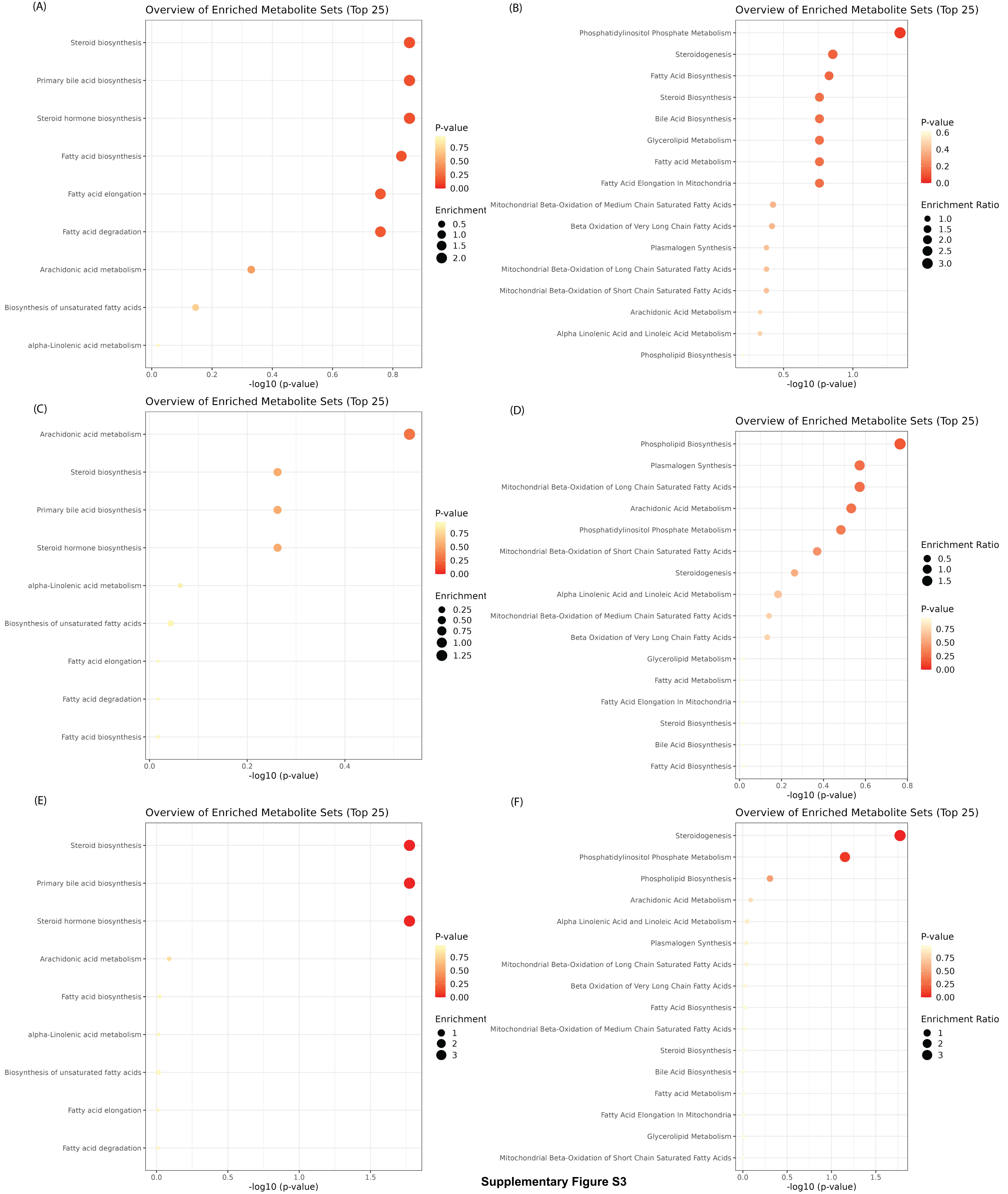


**Supplementary Figure S3**

**Supplementary Figure S3.** Quantitative Pathway Enrichment Analysis. Dot plots representing pathway enrichment analysis across different experimental conditions (Control, R0, and R8) using KEGG and SMPDB databases. Each panel corresponds to a specific comparison, displaying the top 25 enriched pathways. The x-axis represents the statistical significance of enrichment (-log10(p-value)), while the y-axis lists pathway names. The size of each dot represents the enrichment ratio, and the color intensity indicates significance, with darker red indicating stronger statistical significance.

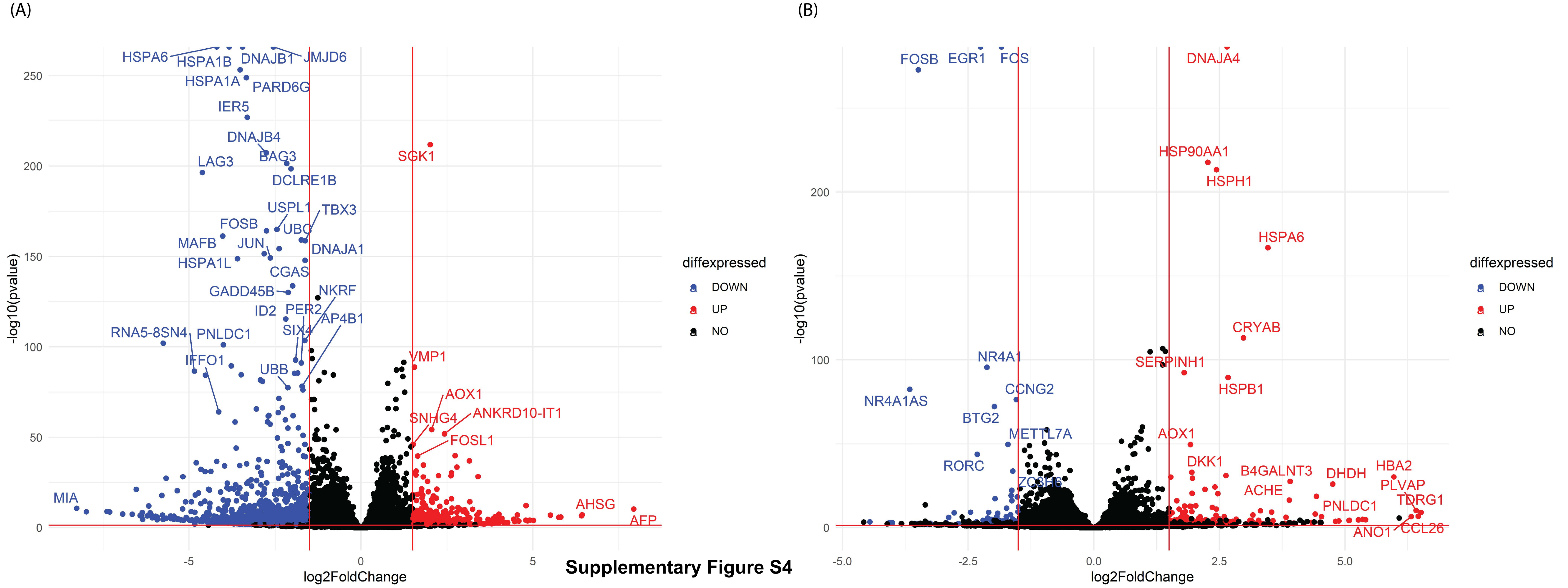


**Supplementary Figure S4**

**Supplementary Figure S4.** Differential Gene Expression in Heat Shock Recovery. (A) Volcano plot illustrating differentially expressed genes in R8 vs. R0. Genes that are significantly upregulated are shown in red, while downregulated genes are shown in blue. (B) Volcano plot depicting gene expression changes in R8 vs. control. Upregulated genes are highlighted in red, downregulated genes in blue, and non-significant genes in black. The figure highlights the widespread transcriptional remodeling occurring during recovery from heat shock.

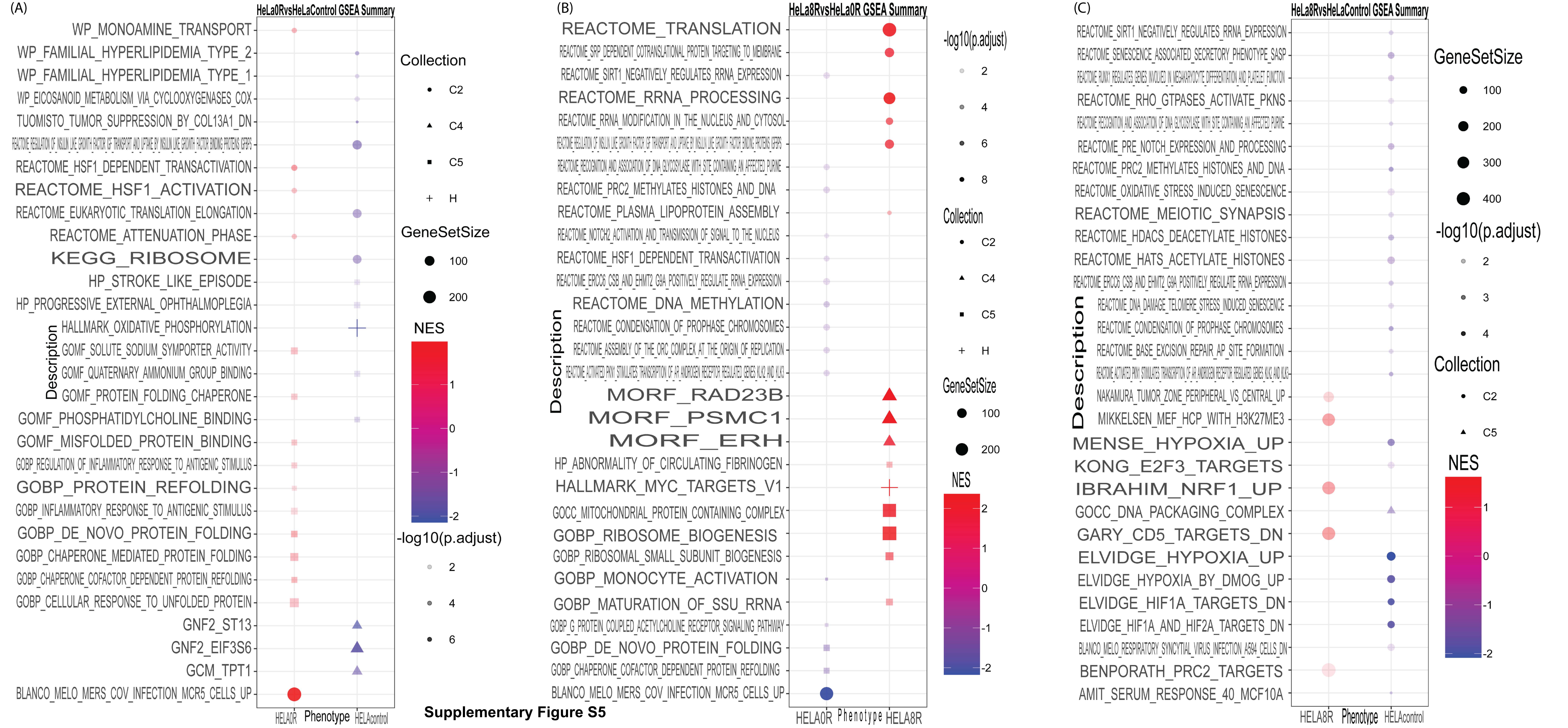


**Supplementary Figure S5**

**Supplementary Figure S5.** Gene Set Enrichment Analysis (GSEA) of Heat Shock Response. (A) Gene set enrichment results for HeLa 0R vs. HeLa control, highlighting enriched pathways related to protein refolding, stress response, and metabolic adaptation. (B) Gene set enrichment results for HeLa 8R vs. HeLa 0R, showing significant enrichment in pathways related to translation, rRNA processing, and chromatin modifications. (C) Gene set enrichment results for HeLa 8R vs. HeLa control, emphasizing pathways associated with transcriptional regulation, histone modifications, and oxidative stress response. Dot size corresponds to gene set size, while color indicates normalized enrichment score (NES), with red representing upregulated pathways and blue indicating downregulated pathways.

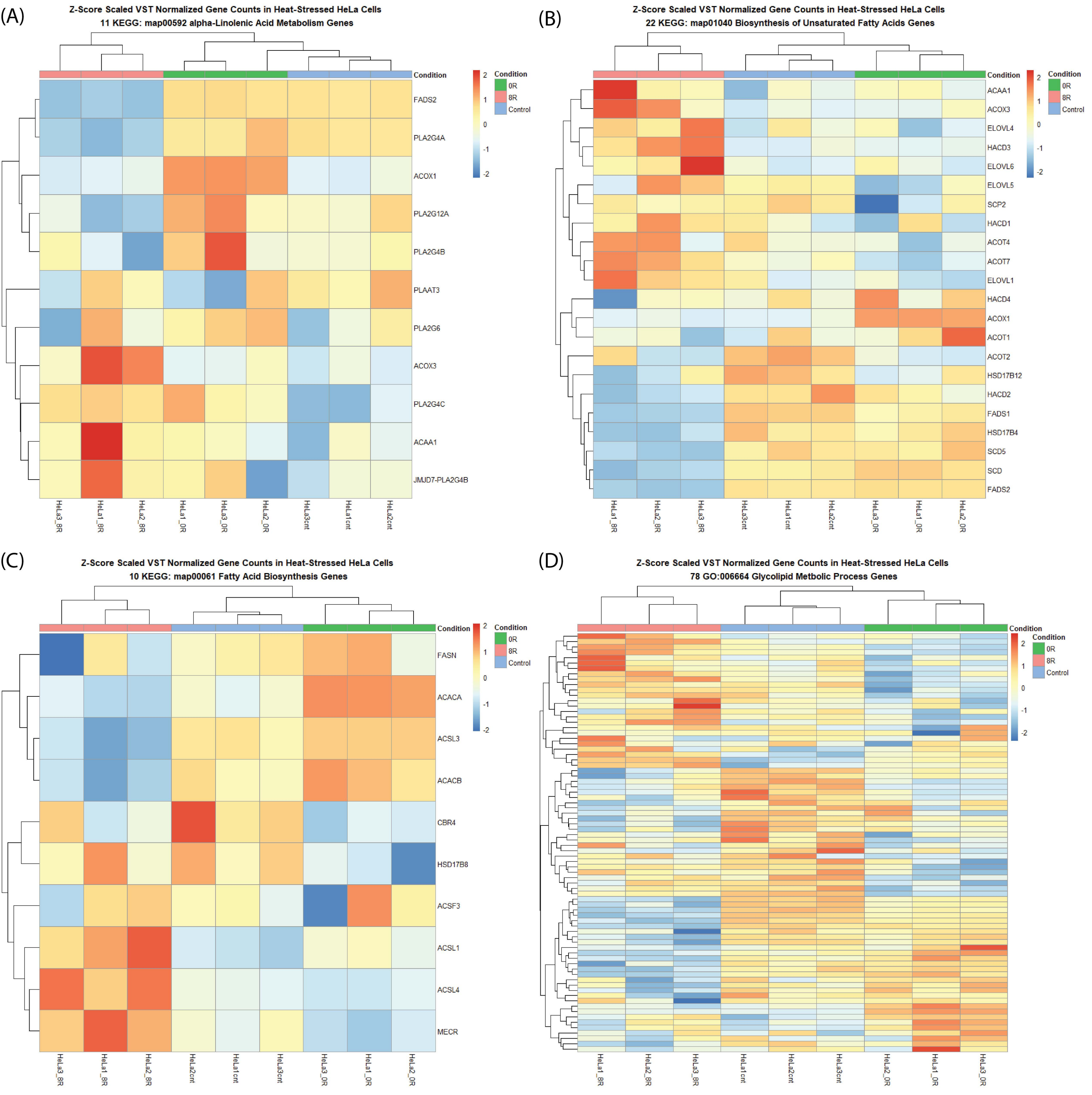


### Supplementary Figure S6

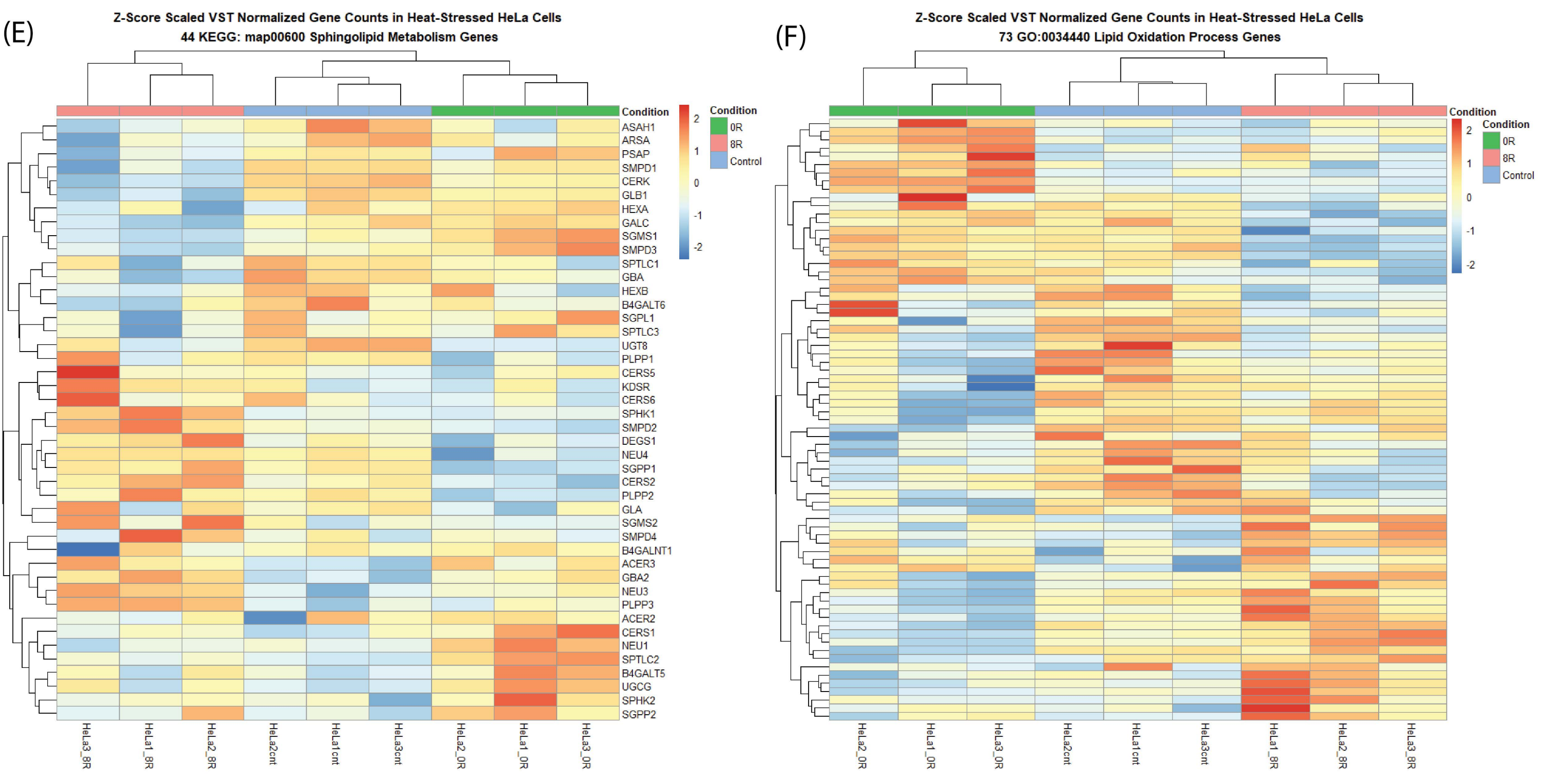

**Supplementary Figure S6.** Heatmaps of Specific Lipid Metabolism Pathways. Z-score normalized gene expression heatmaps for key lipid metabolism pathways across control, R0, and R8 conditions. (A) Shows genes involved in alpha-linolenic acid metabolism, while (B) focuses on biosynthesis of unsaturated fatty acids. (C) Displays fatty acid biosynthesis-related genes, and (D) highlights genes associated with glycolipid metabolic processes. (E) Depicts the expression patterns of sphingolipid metabolism genes, and (F) represents genes involved in lipid oxidation processes. Each heatmap illustrates hierarchical clustering of genes, revealing dynamic transcriptional changes in lipid metabolism in response to heat shock and recovery.

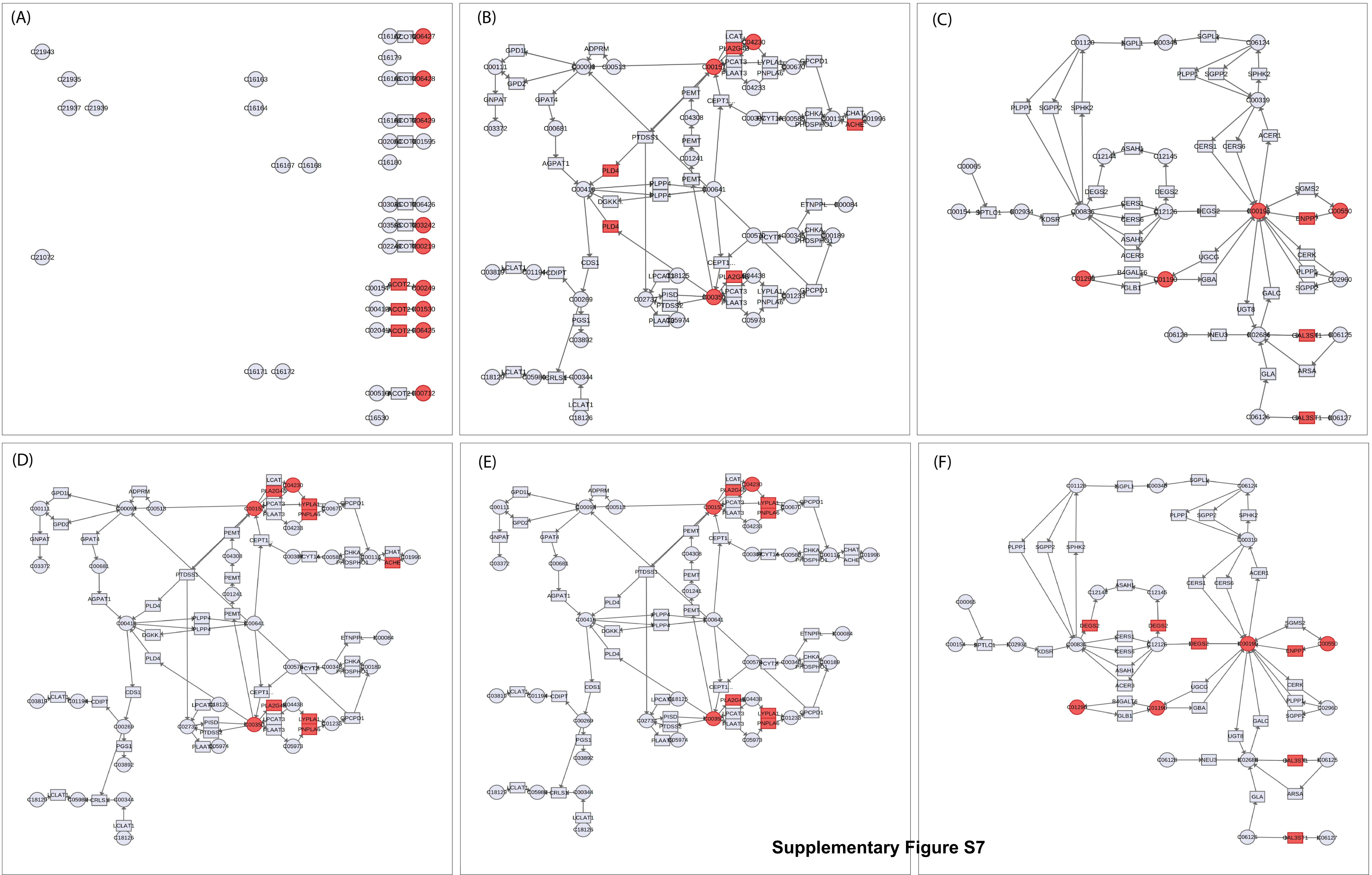


**Supplementary Figure S7**

**Supplementary Figure S7.** Multi-Omics Network Analysis of Heat Shock Response. (A-C) Pathway integration showing gene-lipid interactions in biosynthesis of unsaturated fatty acids, glycerolipid metabolism, and sphingolipid metabolism. (D-F) Pathway integration for recovery (R8 vs. control and R8 vs. R0), highlighting glycerophospholipid metabolism and lipid signaling pathways. Nodes represent genes and lipids, with red-highlighted nodes indicating significant changes in abundance or expression. Edges depict known biochemical interactions and regulatory relationships.
