## Supplementary Table S6 for "Dynamic Lipidome Reorganization in Response to Heat Shock Stress"

**Supplementary Table 6.** Primers used in qPCR experiments

| Accession number | HGNC Symbol | Primer Sequence (5’-> 3’) | Primer Sequence (3’-5’) | Product Size (bp) |
| --- | --- | --- | --- | --- |
| NM_005345.6 | HSPA1A | AGCTGGAGCAGGTGTGTAAC | CAGCAATCTTGGAAAGGCCC | 154 |
| NM_001101.5 | ACTB | CTTCGCGGGCGACGAT | CCACATAGGAATCCTTCTGACC | 104 |
| NM_001256799.3 | GAPDH | CGGGAAGGAAATGAATGGGC | GGAAAAGCATCACCCGGAGG | 148 |
| NM_001114633.2 | PLA2G4B | GGACTCAGTCTCATGGCTGT | GTCAGAGGGGGTCACTAGGTC | 108 |
| NM_006821.6 | ACOT2 | TCATCTCCTGCTGTCCTTCG | TCCGAGCAGGAACCCTAATG | 121 |
| NM_018324.3 | OLAH | TTTTGGCCACAGTATGGGAT | CTTTGGGAATGCGATGCCAG | 140 |
| NM_178543.5 | ENPP7 | TTGTCACCATGACCAGCCC | CCCTCAGGCCCTGCC | 198 |
| NM_003358.3 | UGCG | GTGGCTGATGCATTTCATGGC | CACAAAGGAGCACTTCATATTTGGG | 194 |
| NM_014754.3 | PTDSS1 | TCAGCCTCATGTACTTCGCC | GTCGAGTGAACGGACCATTG | 113 |
| NM_001329544.2 | PTDSS2 | TCCAGACCTCATCCAGCTTAC | GTCAGCGGGGCTACAGTTAG | 135 |
| NM_198580.3 | SLC27A1 | TACATCTACACGTCGGGGAC | GCCGATGATGTTTCCTGCC | 165 |
| NM_013402.7 | FADS1 | GTTCACCCGCCGGCAT | GAACTCATCTGTCAGCTCTTTATTC | 196 |
| NM_001281501.1 | FADS2 | TCACCGGGCAACAGGATG | ACAGGTTCATGTCCTCAGCC | 183 |
| NM_004104.5 | FASN | CTGGAAGGCGGGGCTCTAC | AGTGTGTGTTCCTCGGAGTG | 199 |
